# Menin-MLL1 complex cooperates with NF-Y to promote HCC survival

**DOI:** 10.1101/2025.04.05.647381

**Authors:** Margarita Dzama-Karels, Peyton Kuhlers, Mallory Sokolowski, Jacqueline A. Brinkman, John P. Morris, Jesse R. Raab

**Author notes:** Corresponding author; email:: Contact PI: Jesse R. Raab.

## Abstract

Identification of new therapeutic targets in hepatocellular carcinoma (HCC) remains critical. Chromatin regulating complexes are frequently mutated or aberrantly expressed in HCC, suggesting dysregulation of chromatin environments is a key feature driving liver cancer. To investigate whether the altered chromatin state in HCC cells could be targeted, we designed and utilized an epigenome-focused CRISPR library that targets genes involved in chromatin regulation. This focused approach allowed us to test multiple HCC cell lines in both 2D and 3D growth conditions, which revealed striking differences in the essentiality of genes involved in ubiquitination and multiple chromatin regulators vital for HCC cell survival in 2D but whose loss promoted growth in 3D. We found the core subunits of the menin-MLL1 complex among the strongest essential genes for HCC survival in all screens and thoroughly characterized the mechanism through which the menin-MLL1 complex promotes HCC cell growth. Inhibition of the menin-MLL1 interaction led to global changes in occupancy of the complex with concomitant decreases in H3K4me3 and expression of genes involved in PI3K/AKT/mTOR signaling pathway. Menin inhibition affected chromatin accessibility in HCC cells, revealing that increased chromatin accessibility at sites not bound by menin-MLL1 was associated with the recruitment of the pioneer transcription factor complex NF-Y. A CRISPR/Cas9 screen of chromatin regulators in the presence of menin inhibitor SNDX-5613 revealed a significantly increased cell death when combined with *NFYB* knockout. Together these data show that menin-MLL1 is necessary for HCC cell survival and cooperates with NF-Y to regulate oncogenic gene transcription.

## Introduction

Liver cancer is the third leading cancer-related cause of death worldwide and the most rapidly growing cause of cancer deaths in the United States, with hepatocellular carcinoma (HCC) the most frequent type of liver cancer (∼80%) (1–3). The 5-year relative survival rate of patients with liver cancer combining localized, regional, and distant stages is about 22% (4,5). Curative treatments, including surgical resection, liver transplantation, and local ablation, are available only for early localized HCC. However, most patients are diagnosed at advanced incurable stages of HCC due to challenges in liver cancer screening and late presentation of symptoms, leaving them with systemic therapies and palliative options. Currently, only two single drugs targeting tyrosine kinases (sorafenib and lenvatinib) and three combined immunotherapeutic treatments (atezolizumab/bevacizumab, tremelimumab/durvalumab, and nivolumab/ipilimumab) are approved by the Food and Drug Administration (FDA) as first-line systemic therapies (2). Despite promising advances, most treatment improvements have only resulted in modest survival increases, with many patients remaining unresponsive (6–9). Identifying new vulnerabilities in HCC continues to be a critical medical need.

Aberrant genetic and epigenetic events drive the development and progression of HCC. Changes to the epigenome are proposed to induce cell plasticity and alter the fitness of cancer cells (10). Various epigenetic regulators, including lysine methyltransferases (EZH2, SETDB1, EHMT2), demethylases (KDM3A, KDM4B, KDM5B, KDM1A), and deacetylases (SIRT7, HDAC1-3) were shown to be frequently upregulated in HCC (11–13). Furthermore, sequencing of liver cancer samples has revealed mutations in epigenetic modifiers in about 50% of HCC samples (14–20). Identifying chromatin regulators essential for HCC survival might help develop new targeted therapies to reverse altered chromatin states impacting oncogene expression (11,21,22). Currently, only one epigenetic drug targeting EZH2 has been approved for treating solid tumors (epithelioid sarcoma) despite the clinical success of numerous epigenetic regulator inhibitors in treating hematologic malignancies (12–13), supporting the importance of identifying new epigenetic vulnerabilities in other cancer types. Targeting epigenetic regulators remains a promising area for identifying tumor-specific vulnerabilities (23,24).

We reasoned that the importance of epigenetic regulators in HCC would suggest that further disruption of the epigenome may selectively impact liver cancer cells (25,26). Moreover, focused screens on subsets of the whole genome enable deeper interrogation through the use of multiple models and cell lines. In this study, we used 2D and 3D CRISPR/Cas9 screens of genes involved in chromatin regulation to identify new vulnerabilities in HCC. Several of these targets are core subunits of the menin-MLL1 complex (*MEN1*, *ASH2L*, *KMT2A*). This complex is a current therapeutic target in *MLL*-rearranged (*MLL*- r) and *NPM1-*mutant acute myeloid leukemias (AMLs), where menin inhibitors have shown promising results in Phase I/II clinical trials, and one of them, revumenib (SNDX-5613), being approved by the FDA for treatment of relapsed or refractory *MLL*-r AML treatment (27). MLL1 protein is a ‘writer’ of Histone H3 Lysine 4 trimethylation (H3K4me3), a mark associated with active promoters and has also been suggested to be important in proliferation of HCC and other cancers (28–30). Using a menin inhibitor and CRISPR/Cas9 knockout experiments, we systematically analyzed the global role of the menin-MLL1 complex on HCC survival by performing RNA-seq, CUT&RUN, and ATAC-seq. These data revealed the HCC-specific oncogenic network regulated by the menin-MLL1 complex. We found that active promoters are bound by menin-MLL1 and the transcription factor complex NF-Y. Following menin inhibition, the menin-MLL1 complexes are depleted from chromatin while NF-Y is lost from some targets and re- localized to new sites. Finally, a CRISPR/Cas9 screen of HCC cells in the presence of the menin inhibitor SNDX-5613 revealed potential partners for combined treatments, with the NF-YB transcription factor among the top synergistic hits. We show that the combined inhibition of the menin-MLL1 interaction and *NFYB* gene knockout shows a significantly increased antiproliferative effect in HCC cells. Overall, this study defines the role of the menin-MLL1 complex in coordinating gene regulation to support the proliferation of HCC cells and defines new combinatorial approaches for future HCC therapy improvement.

## Results

### Subunits of the menin-MLL1 complex are a vulnerability in HCC

To identify new potential targets of HCC among epigenetic regulators, we constructed a CRISPR library of ∼6000 guide RNAs (gRNAs) targeting 737 genes involved in chromatin-mediated gene regulation with 7 gRNAs targeting each gene and containing 855 non-targeting controls (Supplemental Table 1). Using this library, we performed an epigenome-focused CRISPR/Cas9 screen on two HCC human cell lines (HLF and PLC/PRF5) in two- (2D) monolayer and three-dimensional (3D) spheroid settings for 28 days (Figure 1A-B, Supplemental Table 2). For 3D conditions, cells were grown on low-adherent plates in methylcellulose (0.75%) containing media to prevent attachment. These conditions have been shown to better mimic *in vivo* growth and revealed distinct vulnerabilities between 2D and 3D CRISPR/Cas9 screens (31). The changes in gRNA representation were calculated relative to the initial timepoint (D0), reflecting the targets that affect the survival and proliferation of HCC cells (Figure 1C-D). A comparison of the 2D and 3D scores revealed a select group of genes essential in both conditions (Figure 1C-D). The gRNAs showing the highest negative scores, indicating the decrease in cell survival upon a gene knockout, were then compared to the Cancer Dependency Map (DepMap24Q4 Public, (32,33) to eliminate commonly essential genes. The common negative hits in all of the performed CRISPR/Cas9 screens included members of the menin-MLL1 complex (*ASH2L, KMT2A, MEN1*), PRC1/PRC2 complexes (*CBX4*, *EED, EZH2, SUZ12*), chromatin remodelers (*ARID1A*, *BAZ2A*, *CHD2*, *SMARCC2*), acetyl- and deacetyl-transferases (*BRPF3*, *CCDC101*, *HDAC7*, *KAT6A*, *KAT7*, *MGEA5*), transcriptional activators or repressors (*RRP8*, *SAFB*, *TADA1*, *TADA3*, *TEAD3*, *TRIM33*) as well as other hits (*ASF1A, BRCA1, GSG2, RAD54L, UHRF1, VRK1*) and represent potential targets for primary HCC (Figure 1E).

**Figure 1.**
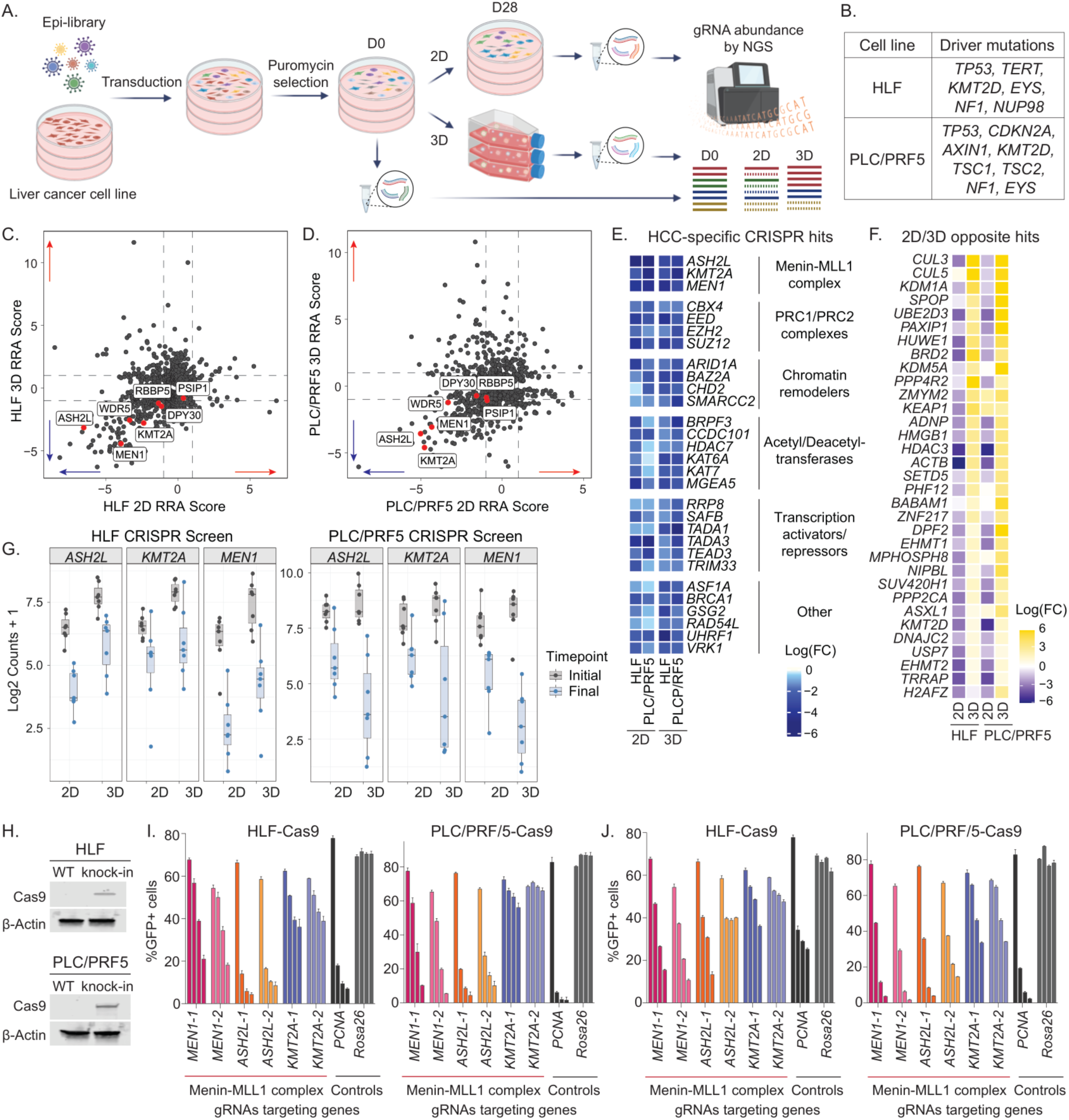
CRISPR/Cas9 screens identify the menin-MLL1 complex as HCC vulnerability. (A) Scheme of epigenome-focused CRISPR/Cas9 screening in 2D and 3D settings. (B) Summary of the most common mutations in HCC cell lines used for CRISPR/Cas9 screening. (C-D) Comparison of 2D and 3D CRISPR screens in HLF (C) and PLC/PRF5 (D) HCC cells using Robust Rank Aggregation (RRA) scores. Subunits of menin-MLL1 complex are shown in red. (E) Heatmap of HCC-specific common negative hits among epigenetic regulators detected by CRISPR/Cas9 screens in HCC cell lines in 2D and 3D settings. Genes are divided in groups based on their functions. (F) Heatmap of hits showing opposite enrichment in 2D and 3D CRISPR/Cas9 screens in HCC cell lines. (G) Logarithmic counts of sequenced sgRNAs targeting *ASH2L*, *MEN1*, or *KMT2A* genes in 2D and 3D screens in HLF (left) and PLC/PRF5 (right) cells comparing initial and final timepoints. (H) Immunoblotting against Cas9 and βActin proteins in HLF and PLC/PRF5 parental and knock-in cells. (I-J) Summary of negative selection CRISPR/Cas9 screen with 2 sgRNAs targeting either *MEN1*, *ASH2L* or *KMT2A* gene or 1 sgRNA targeting either *PCNA* or *Rosa26* serving as positive or negative controls, respectively, in 2D (I) and 3D (J) depicted at d3, d14, d21, d28. Bar graphs represent the mean of measurements of 3 independently transduced cells using the same lentivirus. Error bars represent standard deviation (SD).

For some genes, we observed dramatic differences between 2D and 3D screens in HLF and PLC/PRF5 cells, consistent with recent work in lung adenocarcinoma (Figure 1F, (31)). We detected 33 genes showing opposite enrichment in 2D and 3D screens in HLF cells, including *KEAP1*, a known tumor suppressor that also mediates sorafenib, lenvatinib, and regorafenib resistance (34,35). Multiple genes involved in ubiquitination (*CUL3*, *CUL5*, *UBE2D3*, *HUWE1*, *SPOP*) showed a proliferation benefit for HCC cells only when knocked out in 3D CRISPR screens, suggesting that ubiquitination plays a restrictive role in HCC growth in 3D conditions. Notably, *CUL3*, *UBE2D3*, and *HUWE1* are all considered common essential genes in DepMap. These findings support the importance of CRISPR screening in the 3D spheroid model and highlight the impact of growth conditions on phenotype.

When analyzing the individual gRNA results for members of the menin-MLL1 complex, we confirmed that gRNAs targeting multiple complex members were depleted in both 2D and 3D conditions (Figure 1G). To validate the importance of the menin-MLL1 complex members in the orthologous assay, we performed a CRISPR/Cas9 competitive growth assay using two gRNAs each for *MEN1*, *ASH2L*, and *KMT2A* genes with *PCNA* as positive controls and *Rosa26* as a negative control in Cas9-expressing HLF and PLC/PRF5 cells under both 2D and 3D growth conditions (Figure 1H-J). These data showed a strong dependency of HCC cells on *MEN1* and *ASH2L* genes and less on *KMT2A*, confirming the importance of menin and ASH2L as a part of the menin-MLL1 complex in HCC cell growth. Together, these results suggest that the menin-MLL1 complex is necessary for the survival of hepatocellular carcinoma.

### Menin inhibition strongly affects localization and activity of the menin-MLL1 complex on chromatin

To evaluate HCC cell survival by inhibiting the menin-MLL1 interaction without requiring a complete knockout of the *MEN1* gene, we utilized the publicly available small molecule SNDX-5613 (revumenib). After treating cells for 14 days in 2D (Figure 2A) and 28 days in 3D (Figure 2B-C), we observed a profound dose-dependent cell viability reduction in all tested HCC cells (HLF, PLC/PRF5, SNU398, SNU449), whereas a normal murine liver epithelial cell line, AML12, showed a slight increase in proliferation upon SNDX-5613 treatment. These results support the use of menin inhibitors to dissect the function of the menin-MLL complex in HCC.

**Figure 2.**
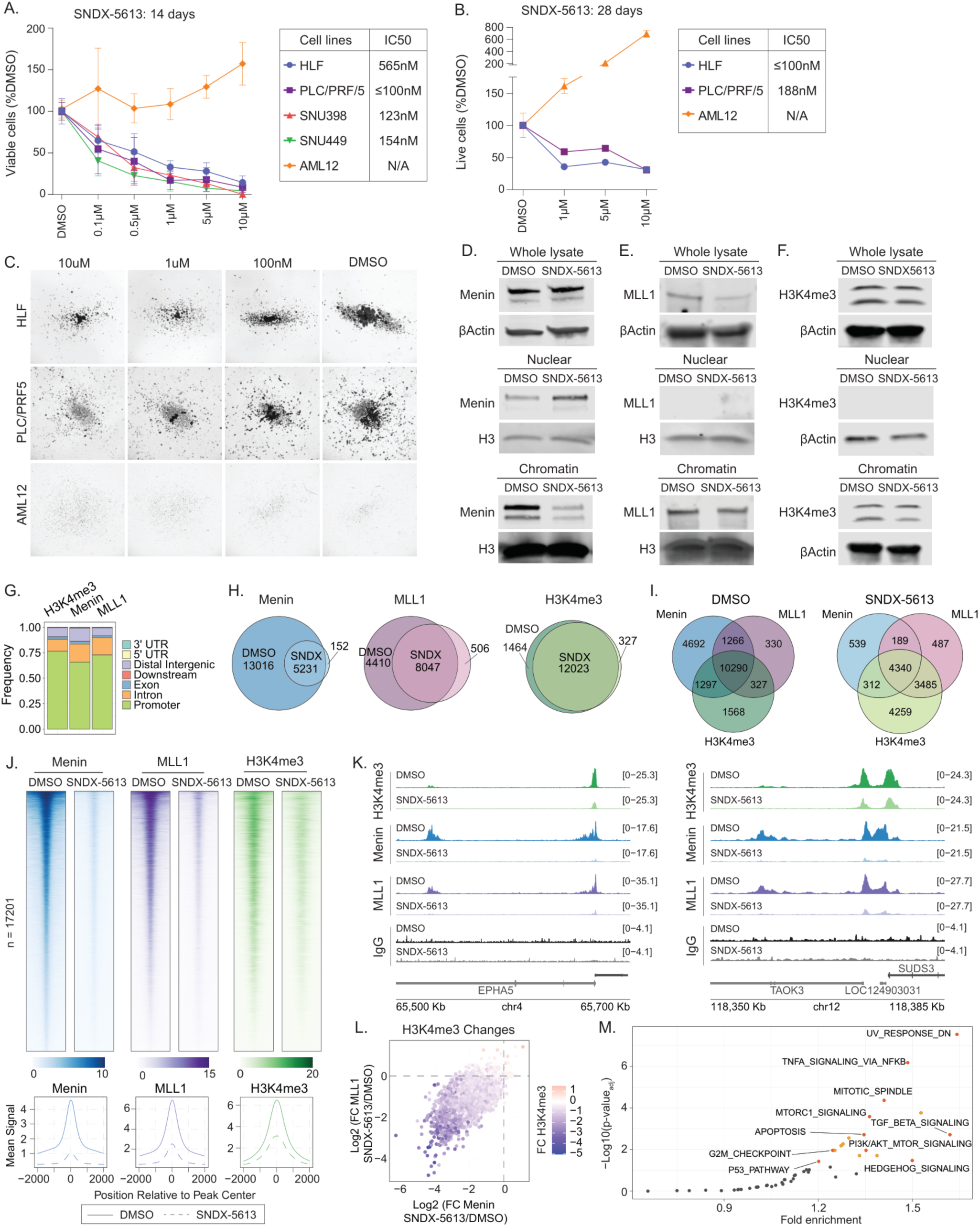
Inhibition of the menin-MLL1 interaction affects HCC cell survival by perturbing the complex binding to set of genes. (A) HCC and murine liver cell viability following SNDX-5613 treatment assessed by Cell Titer Glo (CTG) assay after 14 days in 2D. IC_50_ values were calculated with GraphPad Prism version 9 software. Error bars represent SD of 3 independent experiments, each performed in 4 technical replicates. (B-C) HCC and murine liver cell viability and growth after 28 days of SNDX-5613 treatment with weekly colony dissociation using Accutase and replating in 3D methylcellulose-containing media. Cell viability was measured using CTG assay (B) and cell growth and colony formation ability was assessed by microscopy (C). Micrographs were taken at ×1.25 amplification. (D-F) Immunoblotting against menin (D), MLL1 (E), and H3K4me3 (F) with corresponding controls as βActin or total H3 in HLF cells following treatment with 5µM SNDX-5613 for 4 days (G) Distribution of binding sites relative to genomic features for H3K4me3, menin, and MLL1. (H-I) Venn diagrams of detected independent (H) or overlapping (I) menin, MLL1 and H3K4me3 peaks in HLF cells following treatment with DMSO or 5µM SNDX-5613 (SNDX) for 4 days. Occupancy peaks for all the groups are selected based on menin binding sites. (J) Heatmaps showing the correlation of promoter peaks in a ±2-kb window with occupancy of H3K4me3, menin, and MLL1 across CUT&RUN-seq data from HLF cells treated with 5µM SNDX-5613 for 4 days. Bottom metaplots show the mean of overall peak signals detected at the regions comparing DMSO and SNDX-5613 treatment conditions in HLF cells. (K) Examples of CUT&RUN tracks showing distribution of H3K4me3, menin, MLL1, and IgG binding to the promoter regions in HLF cells treated with 5µM SNDX-5613 for 4 days. The y-axis was chosen for optimal visualization of peaks for each sample. (L) Comparison of logarithmic fold changes (FC) of H3K4me3 mark, menin and MLL1 protein binding to the menin sites comparing conditions of HLF cells treated with 5µM SNDX-5613 to DMSO for 4 days. (M) GREAT analysis of the menin-MLL complex targets using the hallmark gene set as a reference comparing HLF cells treated with 5µM SNDX-5613 to DMSO for 4 days.

Next, we evaluated the impact of menin inhibition on menin-MLL1 complex binding to chromatin. We treated HLF cells with 5µM SNDX-5613 for 4 days, when cells largely remained viable, allowing us to separate the chromatin and transcriptional changes induced by menin inhibition rather than cellular death (Supplemental Figure 1). While total menin protein levels were not affected in HLF cells following treatment with 5µM SNDX-5613 for 4 days, we observed its robust removal from chromatin and accumulation in nuclear fraction as assessed by immunoblotting (Figure 2D). MLL1 protein was also modestly reduced upon SNDX-5613 treatment in both total protein and chromatin fraction, while the global levels of H3K4me3 mark did not show any visible decrease (Figure 2E-F).

Next, we assessed the impact of menin inhibition on the genomic occupancy of menin and MLL1 proteins and the distribution of H3K4me3 using CUT&RUN (36). In HLF cells treated with vehicle, we identified 28123 menin, 17791 MLL1, and 13739 H3K4me3 peaks, with the majority localized to promoter regions (Figure 2G-H, Supplemental Table 3). Comparing peaks in the DMSO and SNDX-5613 treated conditions showed a significant decrease in the menin peak numbers after SNDX-5613 treatment. MLL1 peaks were also reduced, while H3K4me3 peaks were largely unchanged in this analysis (Figure 2H). Notably, the binding sites of menin, MLL1, and H3K4me3 showed significant overlap, consistent with the established role of menin-MLL1 in H3K4me3 methylation at promoters (Figure 2I). Following treatment with 5µM SNDX-5613 for 4 days, we noted a ∼90% reduction in menin signal, a ∼3-fold reduction in MLL1, and a ∼2-fold reduction in H3K4me3 (Figure 2 J-K, Supplemental Figure 2A). Differential occupancy analysis confirmed the overall decrease in menin binding as well as a reduction in the magnitude of MLL1 binding and H3K4me3 (Supplemental Figure 2B). Most sites exhibited a coordinated decrease in menin, MLL1, and H3K4me3 signals, while fewer sites maintained a stable level of H3K4me3 even after the menin- MLL1 complex was removed from the region (Figure 2L, Supplemental Figure 2C). To evaluate pathways regulated by the menin-MLL1 complex in HCC, we performed ontology analysis using the Genomic Regions Annotation Tool (GREAT, (37)) of gene promoters directly bound by both menin and MLL1 proteins and identified enrichment for the TNFα, TGFβ, MTORC1, PI3K/AKT, and Hedgehog signaling pathways (Figure 2M). Together, these data suggest that the menin-MLL1 complexes regulate a large portion of the transcriptionally active regions of the genome that are enriched for genes involved in signaling and cell survival pathways necessary for HCC cell growth.

### Disruption of menin deregulates expression of multiple critical for HCC signaling pathways

To understand how disruption of menin alters gene expression, we performed RNA sequencing (RNA-seq) in HLF and PLC/PRF5 cells treated with 5µM SNDX-5613 for 4 days as in CUT&RUN experiments. In addition, we generated *MEN1* knockout cell lines using CRISPR/Cas9 (Figure 3A, Supplemental Table 4). We observed a strong correlation between the fold changes of DMSO/SNDX-5613 and NTC/Knockout (KO) with more genes downregulated in both cell lines (Figure 3B-C, Supplemental Figure 3A-B). This is consistent with our result showing that menin inhibition leads to widespread decreases in H3K4me3 occupancy at menin-MLL1-occupied promoters, prompting loss of gene activation. Comparing all conditions, we found 63 upregulated and 261 downregulated genes shared among SNDX-5613 treated and *MEN1* knockout cells (Figure 3D). We also observed a larger number of genes impacted by gene knockout relative to menin inhibition. We noted that CRISPR/Cas9 knockouts of menin were generally unstable, often resulting in a quick rebound of expression likely due to a small population of cells not genetically ablated that survived, while *MEN1* knockout led to lethality.

**Figure 3.**
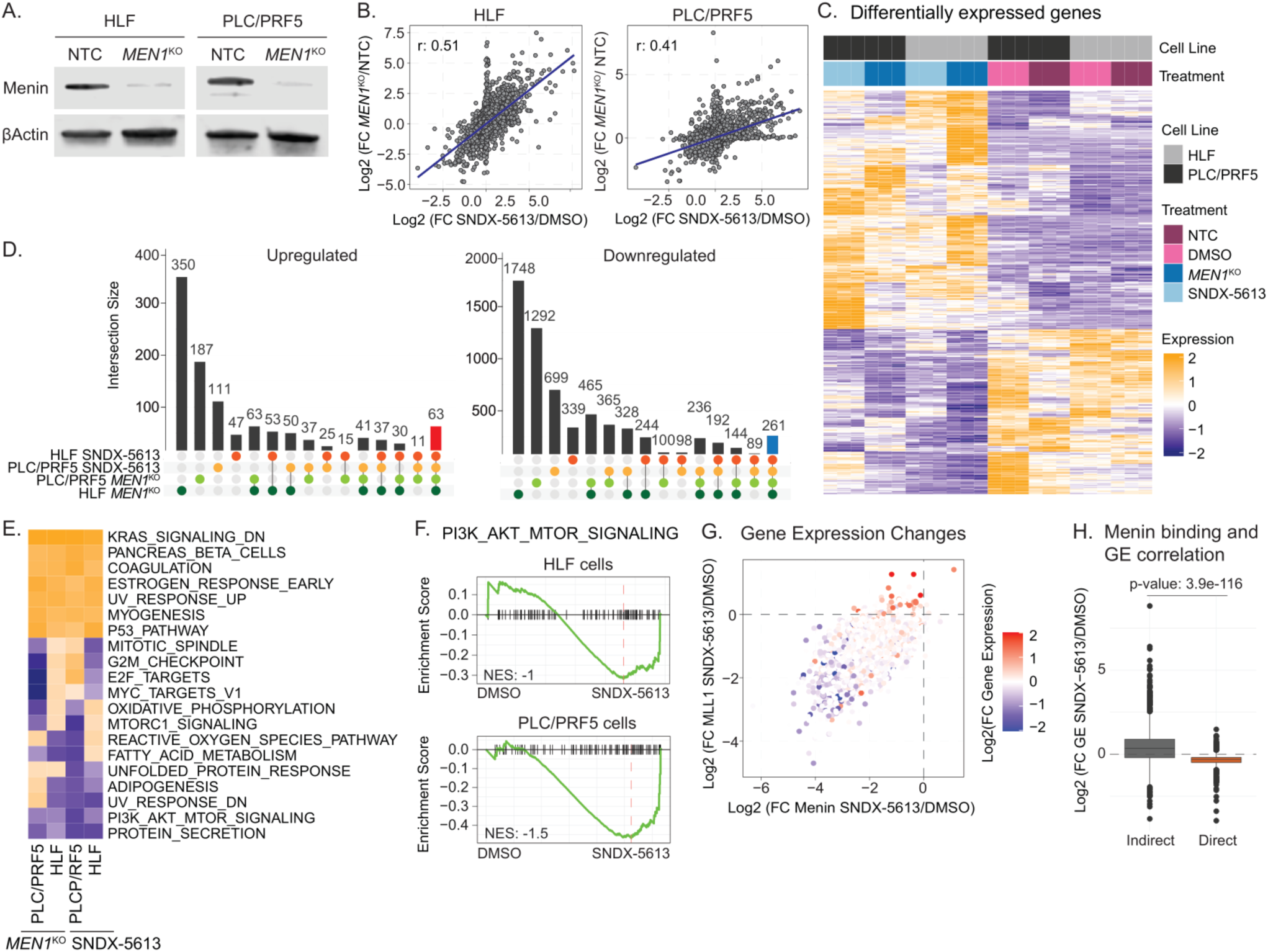
*MEN1* gene knockout and the menin-MLL1 interaction inhibition affects similar signaling pathways in HCC. (A) Immunoblotting against menin and βActin proteins in HLF and PLC/PRF5 knockout (KO) and non-targeting (NTC) cells. (B) Comparison of logarithmic fold changes between *MEN1* knockout relative to NTC and SNDX-5613 treatment relative to DMSO from RNA sequencing of HLF or PLC/PRF5 cells. (C) Heatmap of differentially expressed genes in HLF and PLC/PRF5 cells with *MEN1* or NTC knockout or treated with 5µM SNDX-5613 or DMSO for 4 days. (D) Upset diagrams showing commonly up- and downregulated genes in HLF and PLC/PRF5 cells either treated with 5uM SNDX-5613 or upon *MEN1* knockout. Commonly upregulated genes are marked in red, and commonly downregulated – in blue. (E) Comparison of GSEA ontologies with hallmark gene set used as a reference affected by 5µM SNDX-5613 treatment and *MEN1* knockout in HLF and PLC/PRF5 cells. (F) GSEA analysis of gene expression changes in HLF or PLC/PRF5 cells treated with SNDX-5613 with PI3K/AKT/mTOR signaling pathway from hallmark reference gene set. (G) Comparison of logarithmic fold changes in gene expression and in menin and MLL1 binding to these genes in HLF cells treated with 5µM SNDX-5613 for 4 days. (H) Comparison of logarithmic fold changes in gene expression depending on binding of menin-MLL complex in HLF cells treated with 5µM SNDX-5613 for 4 days.

We then investigated whether menin loss and inhibition led to consistent effects on pathways associated with cancer. We interrogated the MSigDB database using the HALLMARK (H) and Oncogenic Signatures pathway (C6) (38). This revealed a consistent upregulation across both cell lines of EMT, KRAS-associated pathways and Estrogen Response genes (Figure 3D, Supplemental Figure 3C). Downregulated pathways were more variable, but consistent patterns emerged associated with cell cycle, PI3K, AKT, MYC, and YAP (Figure 3E-F, Supplemental Figure 3D). Both MYC and YAP have previously been shown to associate with menin in HCC (39,40).

To investigate the direct impact of menin-MLL1 complex binding on the expression of potential target genes, we integrated CUT&RUN data with the RNA-seq results from HLF cells. We assigned genes as direct menin targets if a menin peak was located within the promoter, menin binding decreased following menin inhibition, and also marked by loss of MLL1 and H3K4me3. This revealed that direct menin targets were downregulated, while indirect targets tended to be upregulated, suggesting the upregulation of pathways we observed in GSEA analysis was likely due to indirect effects of menin inhibition/loss (Figure 3E, G). Directly comparing the correlation between menin and MLL1 binding and expression changes showed that this stringent assignment likely underestimates the number of genes impacted by the disruption of menin and MLL1 binding (Figure 3H). Together, these data suggest the menin-MLL1 complex disruption leads to numerous effects on gene expression and dysregulates key pathways involved in HCC cell survival.

### Menin inhibition causes loss of accessibility at menin-bound regions and alters chromatin accessibility at NF-YB motifs

To understand how inhibition of menin impacts chromatin accessibility, we performed ATAC-seq in HLF cells treated with 5µM SNDX-5613 for 4 days, as in CUT&RUN and RNA-seq experiments. We identified 4236 regions which gained and 2534 regions that lost accessibility (p_adj_ ≤ 0.05, DESeq2) (Figure 4A-B, Supplemental Table 5). Differentially Accessible Regions (DAR) with gained accessibility were associated with promoters, while regions with lost accessibility were more associated with intronic peaks (Supplemental Figure 4A). We further restricted our analysis to the 1565 sites with log2 fold changes greater than 1.5 and 583 sites with log2 fold changes less than 1.5 to determine if changes in accessibility were associated with changes to menin, MLL1, or H3K4me3 occupancy at these DARs (Figure 4C-D). This demonstrated a striking decrease of menin, MLL1, and H3K4me3 binding among the DARs with lost accessibility following menin inhibition. Very few gained sites had evidence of menin or MLL1 binding in either the DMSO or SNDX-5613 treated conditions (Figure 4D). The data support a model where menin promotes open chromatin at transcriptionally active regions and that newly open chromatin sites are independent of menin or MLL1 binding following menin inhibition.

**Figure 4.**
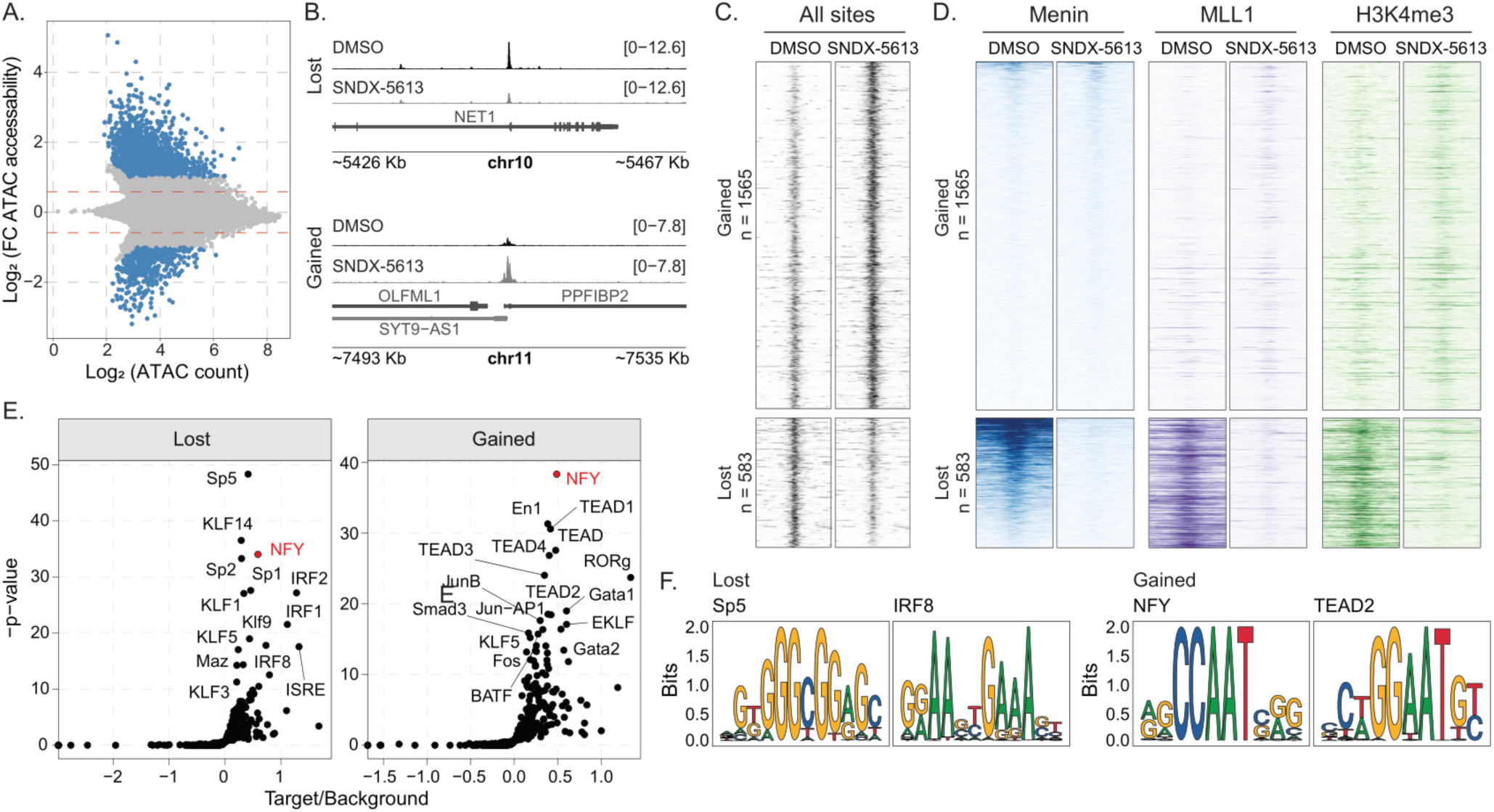
Menin inhibition affects chromatin accessibility in HCC cells. (A) Logarithmic differential chromatin accessibility across ATAC-seq data from HLF cells treated with 5µM SNDX-5613 for 4 days. (B) Representative sequencing tracks of ATAC-seq peaks at the promoter regions with lost and gained chromatin accessibility from HLF cells treated with 5µM SNDX-5613 for 4 days. The y-axis was chosen for optimal visualization of peaks for each sample. (C) Heatmap showing chromatin, H3K4me3, menin, and MLL1 +/- 2kb from peak center at gained or lost ATAC-seq peaks from HLF cells treated with 5µM SNDX-5613 for 4 days (D) Differential binding of transcriptional factors based on motif analysis of gained or lost ATAC-seq peaks relative to unchanged ATAC-seq peaks in HLF cells upon SNDX-5613 treatment (5µM) for 4 days. (E) Examples of motifs derived from the ATAC-seq analysis with lost or gained chromatin accessibility.

To understand which transcription factors may be important for the altered chromatin accessibility at the newly open sites, we used HOMER to search for motifs at gained or lost DARs relative to the stable accessible regions (n = 85708). Lost sites were associated with NF-Y, KLF, SP, and IRF families (Figure 4E-F), while gained sites were also associated with NF-Y, TEAD, and GATA motifs. We observed a similar increase in usage of TEAD and NF-Y motifs using TOBIAS to search for accessibility changes (Supplement Figure 4B). Given the enrichment of NF-Y family motifs in both gained and lost DARs, we hypothesized that NF-Y family members may redistribute in the genome and define a new transcriptional program. NF-Y contains three subunits (NF-YA, NF-YB, and NF-YC), which are all required for its function and have previously been shown to remain bound at menin target sites following menin inhibition and loss from chromatin in *MLL*-r AML (41–43).

### NF-YB relocalizes following menin inhibition

To test if NF-YB changes localization after menin inhibition, we performed CUT&RUN for NF-YB protein following 4 days of treatment with 5µM SNDX-5613. In DMSO treated conditions, ∼40% of NF-YB peaks overlapped with menin peaks. Following treatment with SNDX-5613, the remaining menin sites overlapped ∼77% of NF-YB sites (4178/5388). However, fewer of the NF-YB peaks overlapped with menin peaks (4178/18861, 22%, Figure 5A). By peak overlap we detected 612 NF-YB peaks lost and 2362 peaks gained upon SNDX-5613 treatment (Figure 5B). A more stringent analysis using DESeq2 to identify significantly distinct differentially occupied regions (DORs) revealed 489 lost sites (Lost), 1033 gained sites (Gained), and 18780 sites that were not statistically different (Stable) (Figure 5C). Stable sites were predominately found at promoters, while altered sites were more likely to be found in an intron or distal region (Figure 5D). Notably, about ∼40% of NF-YB sites did not overlap at ATAC-seq peaks in either the DMSO or SNDX-5613 treated conditions (Supplemental Figure 5A). This was unlikely caused by non-specific binding or technical issues in CUT&RUN as the most enriched motif in NF-YB peaks independent of ATAC-seq peaks was the NF-Y motif (Supplemental Figure 5B). These data are also consistent with prior reports that NF-Y could bind closed chromatin or inaccessible regions through a unique histone-like mechanism (43–45).

**Figure 5.**
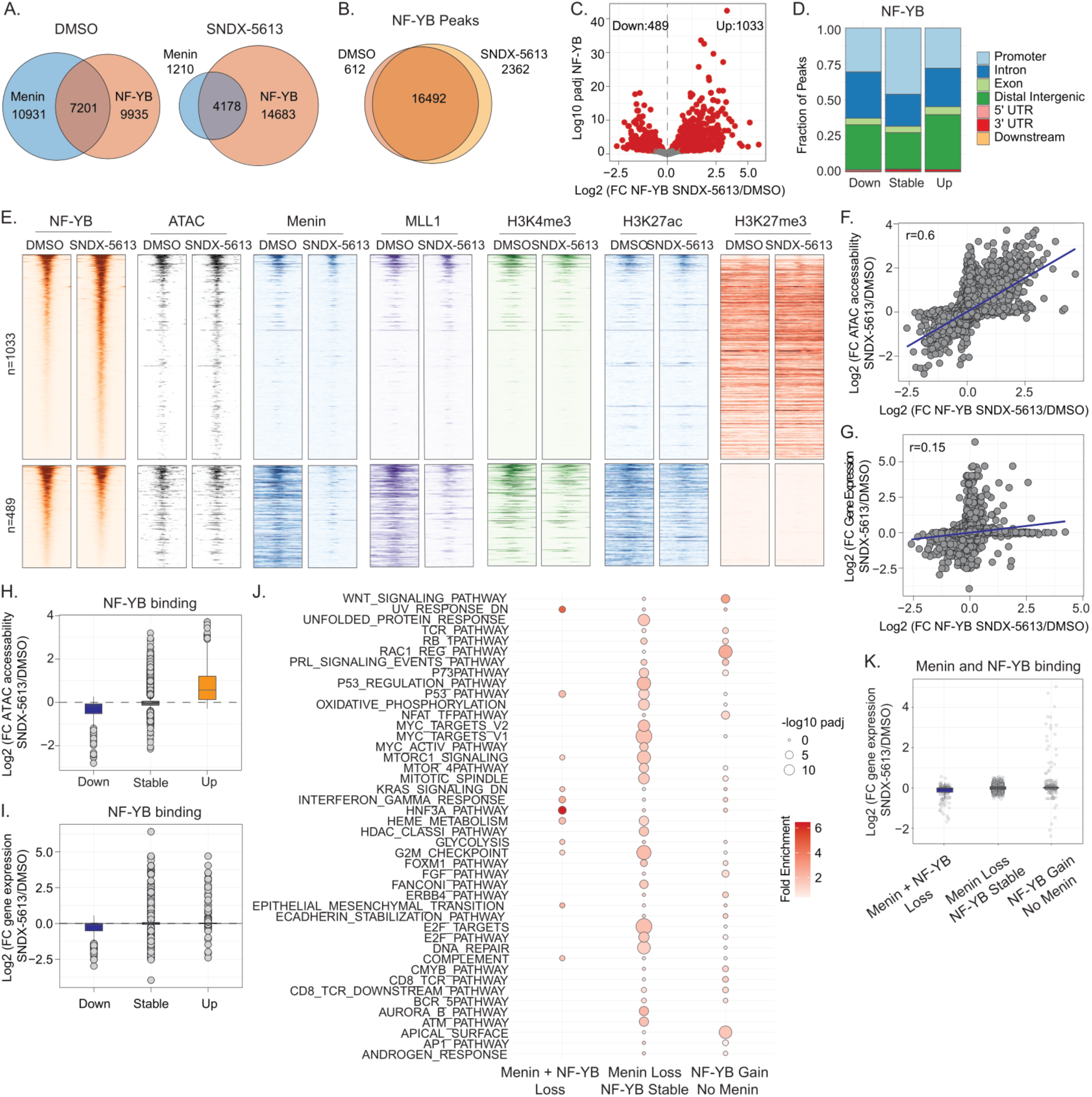
NF-YB binding correlates with newly opened chromatin sites in HCC cells. (A-B) Venn diagram of the overlap between NF-YB peaks and menin peaks (A) or NF-YB peaks alone (B) in HLF cells treated with DMSO or 5µM SNDX-5613 for 4 days. (C) Volcano plot of logarithmic differential binding of NF-YB in HLF cells upon SNDX-5613 treatment (5µM). (D) Distribution of binding sites relative to genomic features for NF-YB based on its binding status (lost, stable, gained). (E) Heatmaps showing the correlation of CUT&RUN or ATAC-seq signal in a ±2-kb window for NF-YB, menin, MLL1, H3K4me3, H3K27ac, H3K27me3 from HLF cells treated with 5µM SNDX-5613 for 4 days at NF-YB gained or lost peaks. (F) Comparison of logarithmic fold changes between NF-YB binding and chromatin accessibility upon SNDX-5613 treatment (5µM) relative to DMSO in HLF cells from CUT&RUN and ATAC-seq data. (G) Comparison of logarithmic fold changes between NF-YB binding and gene expression upon SNDX-5613 treatment (5µM) relative to DMSO in HLF cells from CUT&RUN and RNA-seq data. (H-I) Logarithmic fold change of chromatin accessibility (H) or gene expression (I) based on the NF-YB binding status (lost, stable, gained). (J) GREAT enrichment analysis dot plot of menin and NF-YB direct targets based on their binding status (lost, stable, gained) using the hallmark gene set as a reference comparing HLF cells treated with 5µM SNDX-5613 to DMSO for 4 days. (K) Logarithmic differential gene expression analysis based on menin and NF-YB binding status to corresponding nearest gene in HLF cells treated with 5µM SNDX-5613 to DMSO for 4 days.

Next, we assessed if the loss or gain of NY-YB binding led to additional changes in chromatin accessibility (Figure 5E). We noted that chromatin accessibility was generally decreased at NF-YB lost sites and gained at newly bound NF-YB sites. Similar to our previous finding that menin loss was primarily associated with decreases in accessibility, we found that NF-YB loss was also associated with decreased menin and MLL1 occupancy. Furthermore, NF-YB gained sites did not show a widespread gain in menin or MLL1 binding, suggesting the changes in chromatin accessibility at these sites are not dependent on the menin-MLL complexes. We observed similar changes in H3K4me3 and H3K27ac, further supporting that the primary cause of altered gene expression at sites losing menin and MLL1 is a general decrease in chromatin accessibility associated with combined menin, MLL1, and NF-YB loss leading to an inactive chromatin state.

Our data also revealed that NF-YB gained sites, although associated with increased chromatin accessibility, were still generally embedded in domains of H3K27me3 (Figure 5E). This is consistent with prior studies, which have shown NF-Y binding in both active and repressed chromatin regions (46). These data suggest the sites with gained NF-YB binding may reflect an early activation state that has not fully de-repressed chromatin. Consistent with this, we found a strong correlation between the fold change of NF-YB occupancy and chromatin accessibility (Figure 5F, H, r = 0.6, p-value = 0). However, the expression of the nearest gene to these sites was not as strongly correlated, and most gained NF-YB sites were not associated with a concomitant increase in gene expression (Figure 5G, I, r =0.15, p-value = 7.4e-69). We performed a similar analysis of differential occupancy specifically at accessible chromatin regions (ATAC-seq peaks) and observed that NF-YB binding changes were concordant with chromatin accessibility and, to a lesser extent, with gene expression changes (Figure 5H-I, Supplemental Figure 5C). Together, these data support a model where NF-YB contributes to menin-dependent gene activation, as well as that menin-MLL1 complex and NF-YB are often colocalized, with menin localization being critical for NF-YB localization or stability at these sites. Upon menin inhibition NF-YB relocalizes to novel repressed sites that are poised for future activation, while remaining at some sites in the absence of menin.

To understand the mechanism by which menin inhibition combined with NF-YB loss enhances cell death, we explored our CUT&RUN and RNA-seq data following SNDX-5613 treatment. One possibility consistent with these data is that NF-Y drives a pro-survival transcriptional program following menin inhibition that remain bound by NF-Y and supports continued activation of pathways essential for HCC cell growth. To test this idea, we contrasted regions that either lost both menin and NF-YB, or that lost menin but where NF-YB remained stably bound, or, finally, with NF-YB gained peaks that were not previously associated with a menin peak. We used the Genomic Regions Annotation Tool (GREAT (37) to associate these peaks with genes and determine if distinct gene expression programs were regulated by the different categories (Figure 5J). This analysis revealed regions that lost both NF-YB and menin were associated with cancer pathways such as p53 and KRAS signaling, as well as liver-specific transcription factors such as HNF3A signaling. Regions that lost menin but retained NF-YB had strong enrichment for a wide range of cell growth pathways such as FOXM, MYC, p53, and cell cycle. NF-YB gained regions independent of menin binding were uniquely associated with RAC1 and WNT signaling pathways. The expression of genes that lost both NF-YB and menin was also downregulated after menin inhibition (Figure 5K). The group that lost only menin led to minimal differences in expression (Figure 5K), consistent with our hypothesis that these sites remain active and potentially support cell survival. Finally, there was a modest upregulation of genes that gained NF-YB, but no menin at this early timepoint, suggesting these sites may represent NF-Y priming of associated genes. Together, these data demonstrate that at menin-bound regions NF-YB cooperates to support gene activation and gains new significance by continuing to support the expression of specific pathways and by relocalizing to prime new transcriptional targets.

### NF-YB is a vulnerability in HCC cells following menin inhibition

Next, to identify targets that synthesize HCC cells to menin inhibition, we performed CRISPR/Cas9 epigenome-focused screening in HLF cells treated with SNDX-5613. Following selection to remove non-transduced cells, we treated cells with DMSO or 1µM SNDX-5613 and grew them in 3D conditions for 28 days (Figure 6A). Comparing the gRNA abundance of each condition to their initial distribution revealed several targets that increased sensitivity to menin inhibition. *NFYB* gene knockout was among the strongest SNDX-5613 associated hits consistent with our hypothesis that it re-localizes in the genome following menin inhibition to support the expression of pro-growth genes (Figure 6B-C). To validate these data, we first tested the essentiality of the *NFYB* gene in competitive growth assays. This revealed that NF-YB loss leads to decreased cell viability in HLF cells (Figure 6D). Next, we tested if combined *NFYB* gene knockout and menin inhibition increases HCC cell death by generating two polyclonal CRISPR/Cas9 lines targeting the *NFYB* gene with gRNAs (*NFYB*^KO^, Figure 6E). Treatment of *NFYB*^KO^ and non-targeting control HLF cells with increasing concentrations of SNDX-5613 for 10 days revealed a significantly increased sensitivity to menin inhibition across the entire range of concentrations for both *NFYB*^KO^ cell lines with 500nM of SNDX-5613 being 3X more effective in *NFYB*^KO^ cells (Figure 6F).

**Figure 6.**
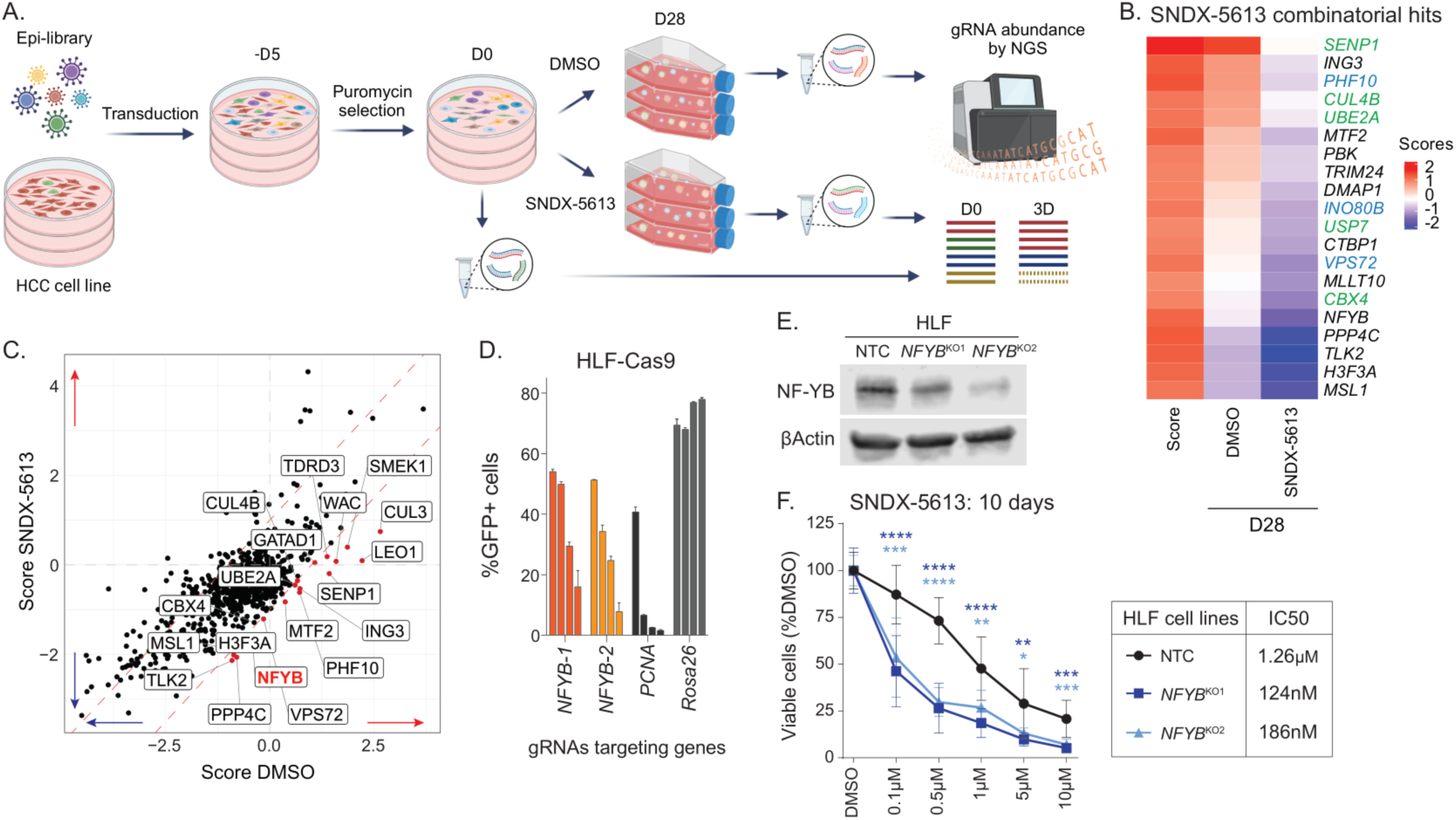
Menin inhibition combined with *NFYB* knockout enhances HCC cell death. (A) Scheme of epigenome-focused CRISPR/Cas9 screening in 3D settings in the presence of SNDX-5613 or DMSO. (B) Heatmap of potential menin inhibitor combinatorial targets from the CRISPR/Cas9 screen performed in HLF cells treated with 1µM SNDX-5613 or DMSO for 28 days. Score was calculated as a substruction of SNDX-5613 from DMSO D28 normalized to D0 scores. Hits representing chromatin remodelers are shown in blue, and hits involved in ubiquitination or SUMOylation – in green. (C) Comparison of gene scores based on gene enrichment in DMSO and SNDX-5613 (1µM) treated groups from 3D CRISPR screens on HLF cells. (D) Summary of negative selection CRISPR/Cas9 screen with 2 sgRNAs targeting *NFYB* gene or 1 sgRNA targeting either *PCNA* or *Rosa26* serving as positive or negative controls, respectively, in 2D depicted at d3, d14, d21, d28. Bar graphs represent the mean of measurements of 3 independently transduced cells using the same lentivirus. Error bars represent SD. (E) Immunoblotting against NF-YB and βActin proteins in HLF knockout (KO) and non-targeting (NTC) cells. (F) Cell viability of HLF cells with either NF-YB knockout or NTC upon SNDX-5613 treatment for 10 days in 2D assessed by CTG assay. IC_50_ values were calculated with GraphPad Prism version 9 software. Error bars represent SD of 3 independent experiments, each performed in 4 technical replicates. P-values were calculated using multiple Student t-test in GraphPad Prism version 9 software. P-values ≤ 0.05 were considered significant with *, P ≤ 0.01 with **, P ≤ 0.001 with ***, P ≤ 0.0001 with ****.

## Discussion

### Expanded use of focused CRISPR screens reveals important differences

In this study, we used epigenome-focused CRISPR libraries to interrogate multiple HCC cell lines in both 2D and 3D culture conditions to test the universality of candidate therapeutic vulnerabilities. Recent work has uncovered that 3D culture conditions can reveal distinct vulnerabilities and more closely mimic essentialities observed *in vivo* (31). While these approaches remain underutilized, our data support that screening approaches should be expanded to include various growth conditions. Notably, we found that the loss of genes involved in protein ubiquitination led to opposing effects on cell growth in 2D and 3D. These genes are often classified as ‘common essential’ in the DepMap database, but when grown in 3D, their loss promotes cell growth (Figure 1). The deficiency of several chromatin regulators detected in our CRISPR/Cas9 screens (*HDAC3*, *KMT2D*, *CUL3*, *CUL5*) have been previously shown rather to promote liver cancer growth, which supports our 3D screening data (47–50). An important logistical consideration when performing CRISPR/Cas9 screens is that smaller pathway-focused libraries can more easily be applied to multiple cell lines and growth conditions, providing an attractive tradeoff compared to genome-wide CRISPR libraries depending on the study’s goals.

### Menin-MLL1 complex has a unique function in gene regulation in HCC

In this study, we showed a dependency of HCC on the core subunits of the menin-MLL histone methyltransferase complex (Figure 1). The interaction between menin and MLL1 was initially found essential in *MLL*-r and *NPM1*-mutant AML subtypes (27,51,52). Subsequent studies have highlighted its significance in the carcinogenesis of solid tumors, such as prostate, breast, Ewing’s sarcoma, endometrial, head and neck, and liver cancer (15). Elevated menin expression in HCC has also been shown to correlate with poor survival prognosis and promotion of HCC development (16). However, the mechanism by which menin-MLL1 promotes HCC previously remained unclear with some reports suggesting *YAP1* or *PEG10* as critical drivers (16–17). With the recent success of clinical trials targeting the menin-MLL1 interaction in AML, understanding menin-MLL complex functioning in HCC could shed light on its potential as an HCC combination therapy.

Our study shows that menin is an important regulator of cell growth in HCC and delineates multiple mechanisms by which it contributes to cancer cell survival. Mechanistically, these differ in some important ways from *MLL*-r and *NPM1*-mutant AMLs, where small molecule inhibitor disruption of the menin-MLL1 interaction partially depletes MLL1 from the genome (53). In HCC, we observe a nearly complete genome-wide depletion of both menin and MLL1 proteins from menin-bound regions with consequent decreases of H3K4me3 and decreased expression at these target genes upon menin inhibition. These data support cancer-type dependent functions of menin-MLL1 complexes in HCC, with MLL1 generally unable to associate with chromatin in the absence of menin (Figure 2). For HCC, we also have not observed menin-MLL1 complex-dependent *MEIS1*- and *PBX3*-driven gene expression found in *MLL*-r and *NPM1*-mutant AMLs. Instead, we identified PI3K/AKT/mTOR signaling as a universally downregulated signaling pathway upon menin inhibition, displaying a completely different mechanism behind the regulation of HCC cell proliferation (27). The significant differences in chromatin accessibility identified in HCC cells after menin inhibition indicated a link between reduced chromatin accessibility at specific sites and the loss of menin-MLL1 complex binding. These sites also showed the strongest decreases in expression (Figure 6H). Notably, we found more sites that gained accessibility following menin inhibition, suggesting that disrupting menin-MLL1 leads to a novel transcriptional response. These sites were largely located away from active promoters defined by H3K4me3 and were not bound by menin or MLL1 in any condition, supporting a hypothesis that these genomic regions represented a menin-MLL1 independent response and potential resistance mechanism in response to menin inhibition treatment.

### NF-Y complex dynamics are driven by inhibiting the menin-MLL1 interaction

We identified the motif for the NF-Y transcription factor complex, shared between NF-YA, NF-YB, and NF-YC, among both gained and lost accessibility regions compared to sites that were stably accessible. NF-Y has previously been shown to localize to the menin-bound regions in *MLL*-r AML (42), where the sites bound by NF-YB following the loss of menin also showed a recruitment of the histone H3 lysine 27 demethylase UTX. While we found sites that could also fit this model, our data suggested two additional mechanisms for NF-Y involvement. First, where the loss of menin could also lead to the loss of NF-Y from those sites, and a second, where NF-Y could bind to novel regions, potentially activating new transcriptional programs. Mapping of NF-Y bound sites following menin inhibition showed NF-Y loss from some menin occupied loci, and these correlated with decreased expression. The sites with gained NF-YB in the absence of menin correlated with moderate increases in accessibility and gene expression (Figure 5), potentially indicating an early response to menin inhibition. While our early time point allowed us to separate the cell death effect from the initial chromatin and transcriptional consequences, it may take longer exposure to menin inhibitors to uncover NF-Y dependent activation. Prior studies of the HCC TCGA cohort found that *NFYA* overexpression in HCC correlates with decreased survival (54). We believe that newly uncovered functions of NF-Y complex upon the menin-MLL1 complex disruption might represent an HCC cell survival mechanism through either supporting previously mobilized oncogenic programs, such as MYC, p53 and E2F targets, or a novel activation of cell invasion and EMT signature genes, usually characterized as escape or metastasis programs (Figure 7, Model).

**Figure 7.**
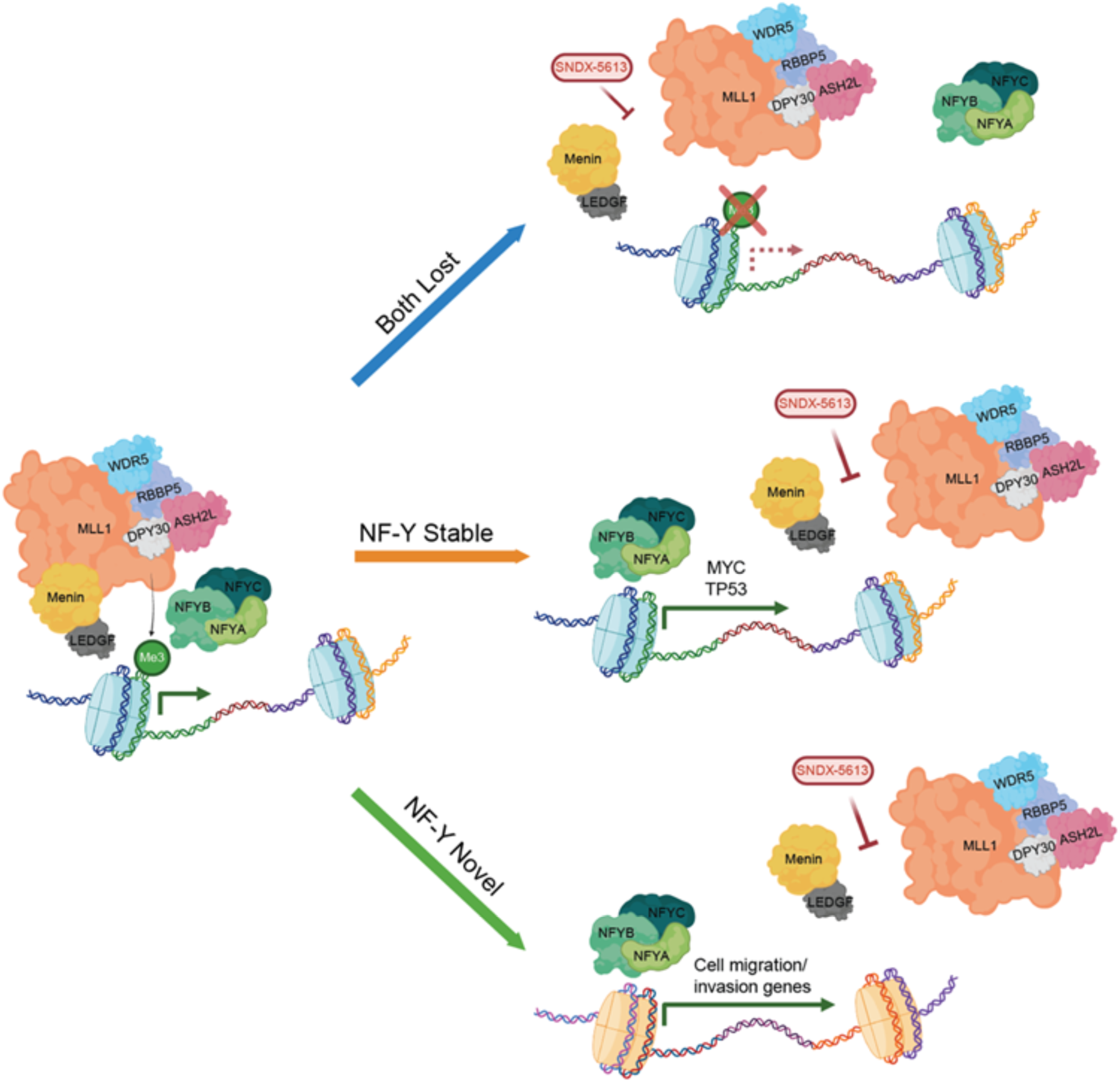
Model of menin-MLL1 and NF-Y complex co-regulation in HCC.

NF-YB was also identified as one of the most promising combinatorial targets for SNDX-5613 treatment in our 3D CRISPR/Cas9 screen, which supports its role in driving survival mechanisms following the disruption of the menin-MLL1 interaction. Furthermore, the treatment of *NFYB* CRISPR/Cas9 knockout cells with SNDX-5613 treatment showed a significant decrease in drug IC50 values, pointing to the necessity of NF-Y complex for HCC cell survival upon menin inhibition (Figure 6). This combined effect may be due to the remainder of NF-Y, which supports continued expression of specific pathways associated with cell growth, such as MYC targets and E2F targets enriched among NF-YB bound regions (Figure 5). Additionally, it could be due to the later activation of novel escape mechanisms depending on NF-Y binding, both trapping cells treated with menin inhibitor in an NF-Y dependent state. Thus, combinatorial treatment represents an appealing evolutionary trap for a potential emerging resistance to menin inhibition in HCC (55). Treatment with menin inhibitors has previously shown emerging resistance in patients carrying a mutation in the *MEN1* gene (56). We believe that the combined menin and NF-YB inhibition could represent a novel route to overcoming menin inhibition resistance. HCC has multiple driver mutations per tumor, which may also contribute to high rates of drug resistance; therefore, finding combinations of vulnerabilities provides an option to improve future therapies (57,58).

Together, our data demonstrate the first comprehensive multi-genomic analysis of menin-MLL1 complex function in HCC and its specific role in HCC cell growth. Here, we demonstrate that NF-Y is a critical modulator of menin-MLL1 function and that both complexes cooperate to maintain an essential HCC survival expression program. The loss of the menin localization at chromatin in HCC cells leads to redistribution of the NF-Y complex and enables novel functions that can be targeted by its disruption in menin inhibited cells, representing a new appealing combinatorial strategy for HCC eradication treatment.

## Methods

### Cell Culture and drug treatment

All HCC (HLF, PLC/PRF5, SNU398, and SNU449) and murine normal liver AML12 cell lines were cultured in a humidified incubator at 37°C with 5% CO2, using the corresponding medias as defined by manufacturer (DMEM, RPMI-1640, or DMEM:F12) supplemented with 10% FBS and 1% Penicillin-Streptomycin (Gibco), and, additionally for AML12 cells, 1x ITS Universal Culture Supplement (Corning). Trypan-excluding cells were plated at a density of 5,000-20,000 cells depending on a cell line per well in 1 ml media in 24 well plates. Cells were split and replated in fresh media every 3-4 days if they were treated for longer than 4 days. For IC50 determination, cells were treated with small molecule inhibitor SNDX-5613 at 10µM, 5µM, 1µM, 500nM, and 100nM concentrations plus vehicle control (DMSO). Cell viability was assessed by CellTiter-Glo® (CTG) luminescent cell viability assay (Promega) using CTG reagent for staining in a ratio with cells 1:1 for 10 min at RT with gentle agitation. The signal was assessed using fluorescence assay on Cytation 5 machine (BioTek).

### Immunoblotting

HCC cells treated with SNDX-5613 compound and vehicle (DMSO) were lysed in RIPA buffer (1e6 cells/30 µl lysis buffer). Identical amounts of protein of each sample were loaded on a 4-15% TGX precast gels (BioRad), electrophoretically separated and transferred to PVDF membranes (BioRad) using a Trans-Blot Turbo machine (BioRad) according to the manufacturer’s protocol. The membranes were blocked with 1:1 ratio mix of Intercept blocking (LI-COR) and TBS (200 mM Tris, 1500 mM NaCl) buffers for 1h at RT. Membranes were cut to allow independent incubation with primary antibodies according to the manufacturer’s recommendations in the antibody mix with ratio 1:1:0.01 of Intercept blocking buffer, TBS buffer, and 10% Tween 20 overnight at 4°C with gentle agitation. Membrane then was washed 3 times with TBST buffer (TBS with 0.1% Tween 20) for 10 min at RT with gentle agitation. The membrane then was incubated with secondary antibodies according to the manufacturer’s recommendations in the antibody mix with ratio 1:2:0.01:0.002 of Intercept blocking buffer, TBS, 10% Tween 20, and 10% SDS for 1h at RT with gentle agitation. The membrane was developed using Odyssey CLx machine (LI-COR). The immunoblotting signal analysis was performed in ImageStudio (LI-COR).

### CRISPR Screen

CRISPR libraries were designed using the GUIDES web app (http://guides.sanjanalab.org/) using a list of genes collated by hand (Supplemental Table 1). Oligos were synthesized by Twist and cloned into pLenticrsipr-V2 as described (59,60). The lentivirus of gRNA library was made by transfecting HEK293T and 2:2:1 ratio of psPAX2:pLenticrispr:pMD2.6 before harvesting viral supernatant at 72h. Cells for screen were infected at a multiplicity of infection of 0.25-0.3, selected for 7 days with 1µg/ml puromycin for non-infected cells. Cell pellets from ≥5e6 cells were harvested for genomic DNA (gDNA) using Monarch Genomic DNA Purification Kit. At least 1.5e6-3e6 cells was then placed under selective pressure (2D or 3D growth, or with 1µM SNDX-5613) for 28 days. This represents > 500X coverage of the CRISPR library. After 28 days of growth, gDNA was harvested as before and high-throughput sequencing libraries were generated by PCR with oligos used for adding Illumina adapters and barcodes. Libraries were sequenced 1x75bp on Nextseq 500.

### CRISPR Screen Analysis

CRISPR/Cas9 screen data were processed using the MaGeCK pipeline(61) using the ***n***on-targeting gRNAs as controls. Robust Ranking Algorithm (RRA scores) from each contrast (2D vs Time 0; 3D vs Time 0) in both cell lines were compared using R. For comparison of individual gRNAs counts were normalized for sequencing depth.

### Competitive Growth Assays

gRNAs targeting genes of interest and control genes (PCNA and Rosa26) were cloned into GFP-gRNA-coupled Lenti_sgRNA_EFS_GFP (LRG) plasmid (Addgene #65656). Lentivirus was made from these plasmids as described above and used to infect HCC cell lines of interest such that ∼50% of initial cells were infected. Percent of GFP+ cells was measured for up to 28 days and compared to GFP+ cells with either a Rosa26 (non-targeting control) or PCNA (essential gene control) gRNA insertions.

### CUT&RUN

CUT&RUN was adapted from (62) and Epicypher. 200,000-500,000 (depending on antibody) cells were pelleted for 3 minutes at 300g at room temperature (RT). Cells were washed twice in wash buffer (20mM HEPES pH 7.6, 150mM NaCl, 0.5mM Spermidine, protease inhibitor (1 tablet/30mL Roche). 10µL Concanavalin A beads (Bangs Labs) per sample were washed twice in Bead Activation Buffer (20mM HEPES ph7.9, 10mM KCl, 1mM CaCl_2_, and 1mM MnCl_2_). Cells were incubated with Concanavalin A beads in Wash buffer for 5-10 minutes at RT. Supernatant was removed and bead bound cells were resuspended in 50µL Wash buffer containing 0.05% Digitonin and 2mM EDTA. Antibodies (See table) were added to mixture and mixed by gentle flicking before incubation overnight at 4°C. Bead bound cells were washed on a magnet 2 times using Wash buffer containing 0.05% Digitonin (Dig-Wash Buffer) before incubation in 50µL Dig-Wash buffer with guinea pig anti-rabbit secondary antibody (1:100, Novus NBP1-72763) for 1 hour at 4°C. Bead bound cells were washed as described above before incubation in 700ng/µL pAG-MNase (purified in-house per (63)) for 1 hour at 4°C. Bead bound cells were washed 4 times as above with cold Dig-Wash buffer then resuspended in 50µL cold Dig-wash buffer while kept on ice. 1µL of 100mM CaCl2 was added while on ice and cells were incubated on ice at 4°C for 30 minutes. Reaction was then quenched by addition of Stop buffer (340mM NaCl, 20mM EDTA, 4mM EGTA, 0.05µg/µL RNAseA (ThermoFisher ), and 0.1% Triton-X100). Cells were incubated at 37°C for 30 minutes in PCR machine before purifying DNA using a Zymo DNA Clean and Concentrator-25. Libraries were constructed using the Kapa Hyperprep Kit with modifications. End-repair and A-Tailing reactions were incubated at 12°C for 15 minutes, 37°C for 15 minutes, and 58°C for 45 Minute. Adapters were ligated for 1 hour using 5µL of 750nM adapter concentration (Roche Dual Index). Libraries were cleaned twice using 1.1X volumes Kapa pure beads before PCR amplification using Kapa Hifi PCR mix with the following conditions: 98°C for 30” to denature then 14 cycles of 98°C for 15”, 60°C for 10” before a final Kapa Pure Bead clean-up using 1.2X bead volumes. Libraries were quantified and pooled before sequencing at 2x50bp or 2x150bp on an Illumina Nextseq 1000 or NovaSeq X plus10B.

### CUT&RUN Analysis

CUT&RUN data was first processed using our CUT&RUN Nextflow pipeline (github.com/raab-lab/cut-n-run Version 4.0) which implements alignment, filtering, sorting, peak calling, and creates bigwig files for visualization. Trimming was performed using trim_galore with default parameters. Alignment was performed using Bowtie2 –very-sensitive-local -X 800 parameters to hg38. Files were sorted and indexed using Samtools and duplicates were marked with Picard. Macs2 was used for peak calling ( --call-summits qvalue=0.05). Consensus peak calls across replicates were generated using the RMPSC package (64). Normalization for visualization was performed using an implementation of the “composition” method of CSAW to calculate normalization factors for each antibody before scaling individual tracks using this factor. Coverage tracks were generated using Deeptools (version 3.2.0). Final visualization was generated by averaging bigwig signal tracks across replicates. These steps are all implemented within the Nextflow pipeline raab-lab/cut-n-run. Downstream analysis was performed in R. Peaks were annotated to the nearest gene and genomic features using ChIPpeakAnno package. Heatmaps and metaplots of CUT&RUN signal were calculated using Deeptools computeMatrix and plotHeatmap functions. Motif analysis was performed using HOMER (65) and TOBIAS (66).

### RNA-seq

1e6 cells were harvested in biological replicates and pelleted for 1 minute at 500g at RT. Total RNA was isolated using the Monarch Total RNA Miniprep Kit including on column DNAse I digestion. 750 ng of total RNA was used for mRNA library preparation using KAPA mRNA HyperPrep Kit and in accordance with manufacturer’s instructions. The desired mean library size was selected in the range of 200-300 bp with fragmentation step for 6 min at 94°C. For adapter ligation, 7µM adapter stock was used, and 9 cycles were chosen for library amplification. Libraries were quantified and pooled before sequencing at 1x75bp on an Illumina Nextseq 1500.

### RNA-seq Analysis

FASTQ files were processed using our Nextflow pipeline (github.com/raab-lab/rnaseq) to generate count matrices for each transcript using Salmon (67). Differential expression analysis was performed by first importing and merging transcripts to genes using tximeta before DESeq2 to perform differential expression. Log2FoldChanges were shrunk using the lfcShrink(type = ‘apeglm’) function. All downstream plotting was performed using R. GSEA analysis was performed using clusterProfiler (version = 4.10.1) with msidgdb (38) to provide gene annotations.

### ATAC-seq

ATAC-seq was performed using the Omni-ATAC protocol. Briefly, 50,000 cells were pelleted at 500gx5 minutes at 4°C. Cells were resuspended in 50uL of cold ATAC-RSB (10 mM Tris-HCl pH 7.4, 10 mM NaCl, and 3 mM MgCl_2_ in water) containing 0.1% IPEGAL-CA630, 0.1% Tween-20, and 0.01% Digitonin and pipetted up and down 3 times before incubating on ice for 3 minutes. The Lysis buffer was then washed out using 1mL cold ATAC-RSB containing 0.1% Tween-20 but no IPEGAL-CA630 or Digitonin. Nuclei were then pelleted at 500g x10 minutes at 4°C. All supernatant was removed, and pellet was resuspended in 50uL of transposition mix - TD buffer, 0.05% Digitonin, 0.1% Tween, and 200nM TN5 transposase (Diagenode). Cells were incubated at 37°C for 30 minutes on a Thermomixer at 1000RPM. Reactions were cleaned using a Zymo DNA Clean and Concentrator-5 (D4014, Zymo Research). Libraries were amplified to add indexed adapters from under the following conditions. 72°C x 5 minutes, 98°Cx30” followed by 5 cycles of 98°Cx10”, 63°Cx30”, 72°Cx1’. The amount of library was quantified using qPCR to select the correct total number of cycles as described in (68), before completing cycle as before. Final library was purified using a 2-sided bead clean up (Roche Kapa Pure Beads), with ratios of 0.6X (keeping the supernatant) and 1.2X (keeping the bound fraction) to remove large fragments and unincorporated adapters respectively. Libraries were then pooled and sequenced on a Nextseq 1000 (Illumina) 2x50bp.

### ATAC-seq Analysis

Fastq files were processed using our Nextlfow pipeline (github.com/raab-lab/cut-n-run –atac). This pipeline is similar to our CUT&RUN approach, with some ATAC-seq specific changes denoted by the –atac flag. Briefly, reads are aligned with Bowtie2, duplicates are marked with PICARD, normalization factors are calculated using CSAW ‘efficiency’ approach on a consensus set of peaks from all replicates/conditions defined using RMSPC, as for CUT&RUN. For differential accessibility, reads overlapping consensus peaks were counted using the CSAW function regionCounts and then used as input for DESeq2 to perform differential testing and lfcShrink(type = ‘apeglm’). Annotations to nearest feature and gene were performed using ChIPpeakAnno (version).

## Statistical Analysis

All computational analysis of genomics data was performed using the R programming language (version 4.3.2) unless otherwise noted. Drug sensitivity and cell growth assays were analyzed in Prism GraphPad (version = 9).

## Reagents

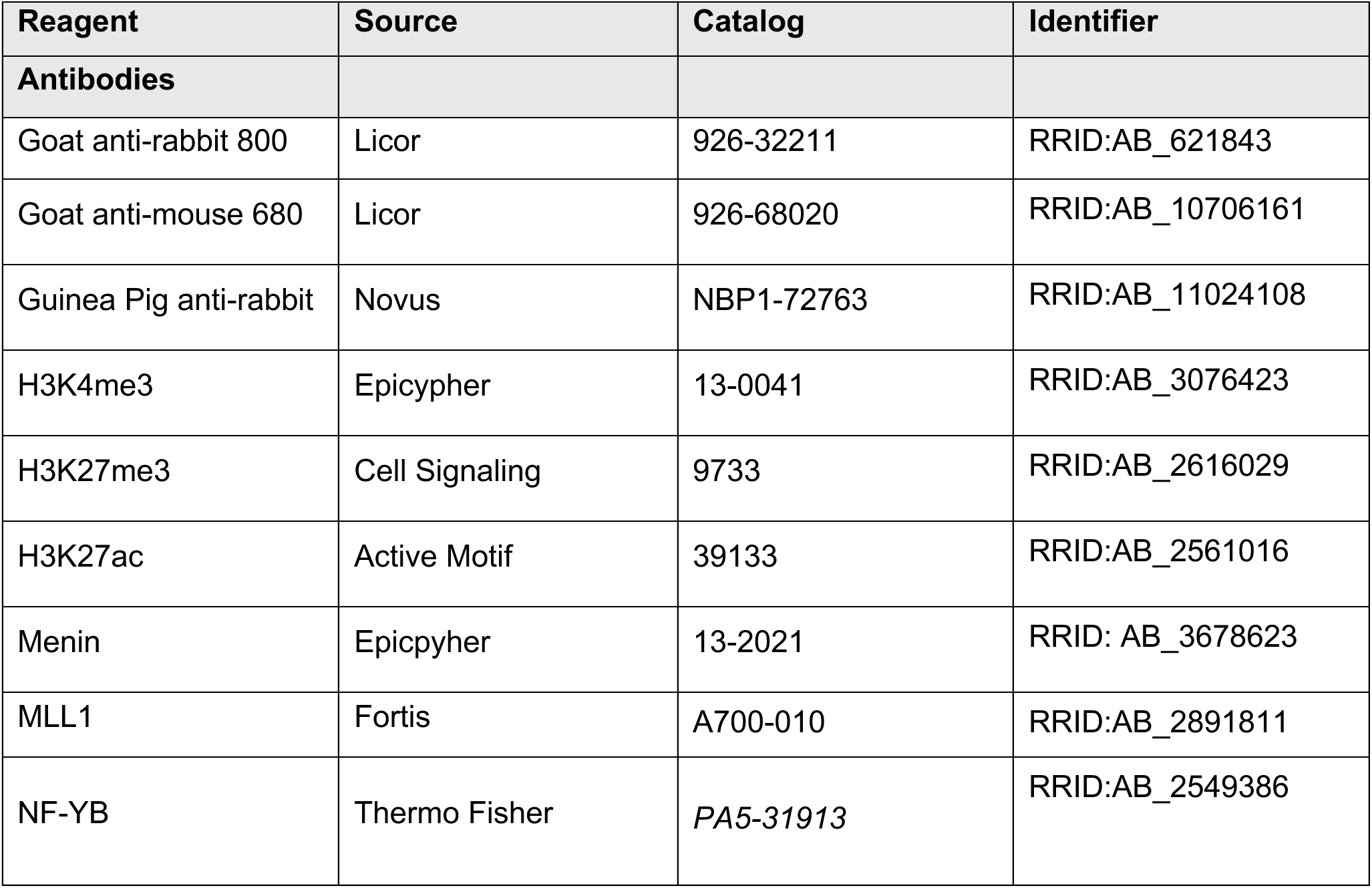
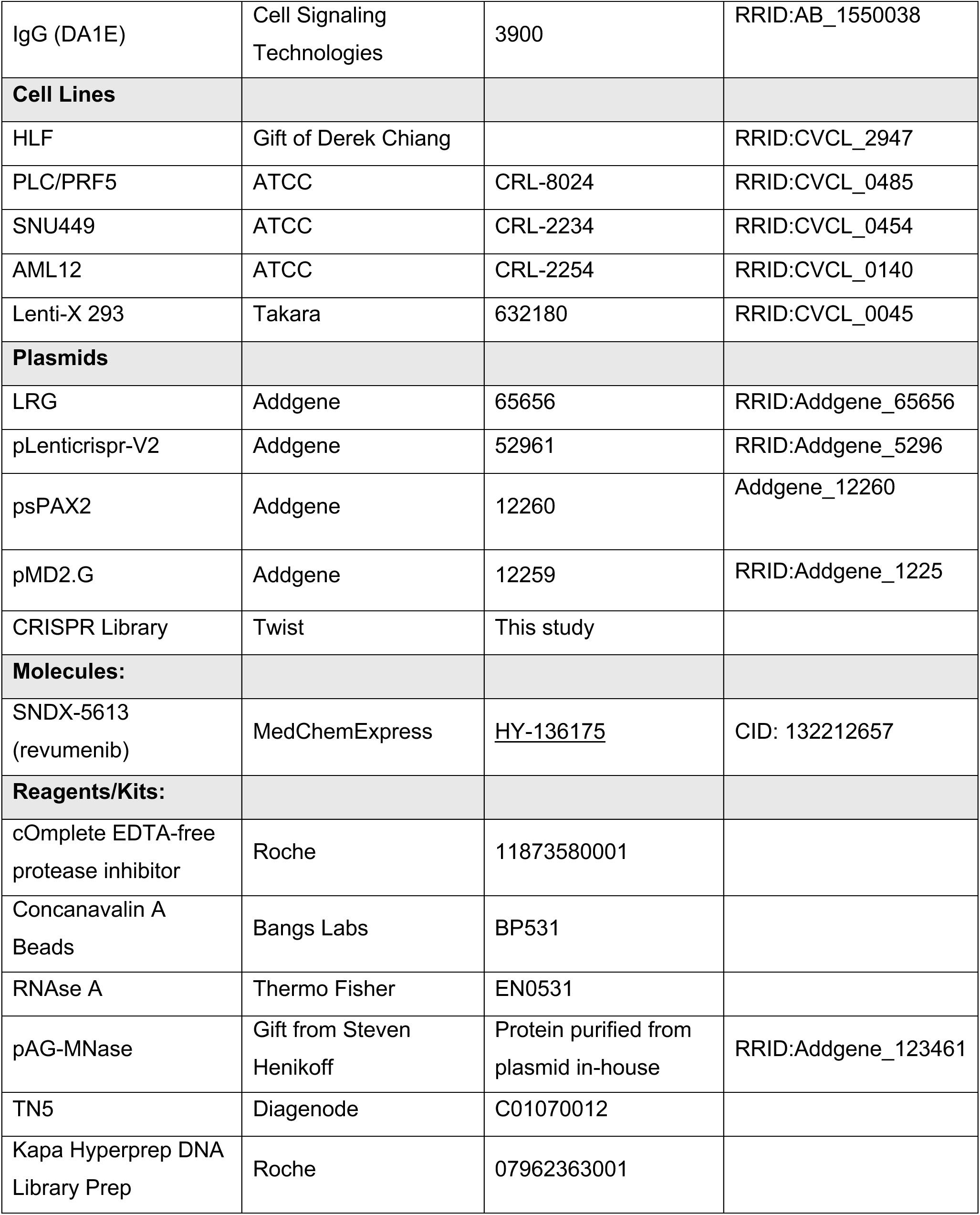
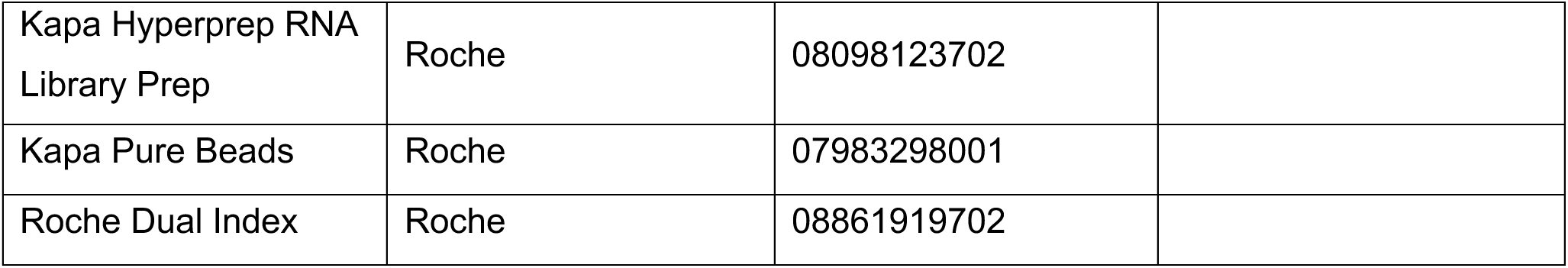

## Code and Data Availability

Raw and processed sequencing data for CUT&RUN, RNA-seq, and ATAC-seq will be deposited at GEO at acceptance. Processed data for fold changes and p-values for each experiment are included as Supplemental Tables. CRISPR screen RRA scores are found in Supplemental Table 2. Code for processing CRISPR, RNA-seq, CUT&RUN, and ATAC are available at github.com/raab-lab/ and code used for analysis of data at github.com/raab-lab/dzama2025-menin-mll

## Author Contributions

MDK and JRR conceived of the study. MDK, JRR, and JPM designed experiments. MDK, MS, and JRR performed experiments. MDK and JRR wrote the manuscript. MDK, PK, JAB, and JRR performed data analysis. All authors edited and approved of the manuscript.

## Acknowledgements

We thank members of the Raab Lab for their helpful comments. This study was supported by internal support from the Lineberger Comprehensive Cancer Center to JRR. MD was supported by ACS Postdoctoral Fellowship (PF-24-1247688-01-DMC). MS was supported by T32GM135128. JAB was supported by R35GM147286-02S1 pAG/MNase was a gift from Steven Henikoff (Addgene plasmid #123461; http://n2t.net/addgene:123461; RRID:Addgene_123461). LRG (Lenti_sgRNA_EFS_GFP) was a gift from Christopher Vakoc (Addgene plasmid# 65656; http://n2t.net/addgene:65656; RRID:Addgene_65656). Plasmid Lenticrispr-V2 lentiCRISPR v2 was a gift from Feng Zhang (Addgene plasmid #52961; http://n2t.net/addgene:52961; RRID:Addgene 52961), psPAX2, pMD2.G.

---

**Supplemental Figure 1.**
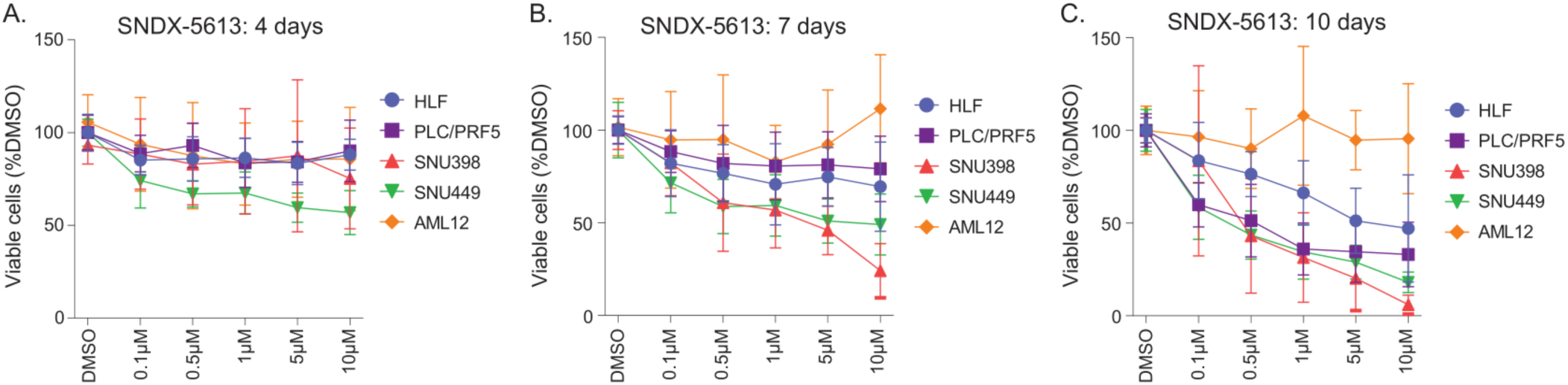
Inhibition of the menin-MLL1 interaction affects HCC cell survival. (A-C) HCC and murine liver cell viability following SNDX-5613 treatment assessed by Cell Titer Glo (CTG) assay after 4 (A), 7 (B) or 10 (C) days in 2D. Error bars represent SD of 3 independent experiments, each performed in 4 technical replicates.

**Supplemental Figure 2.**
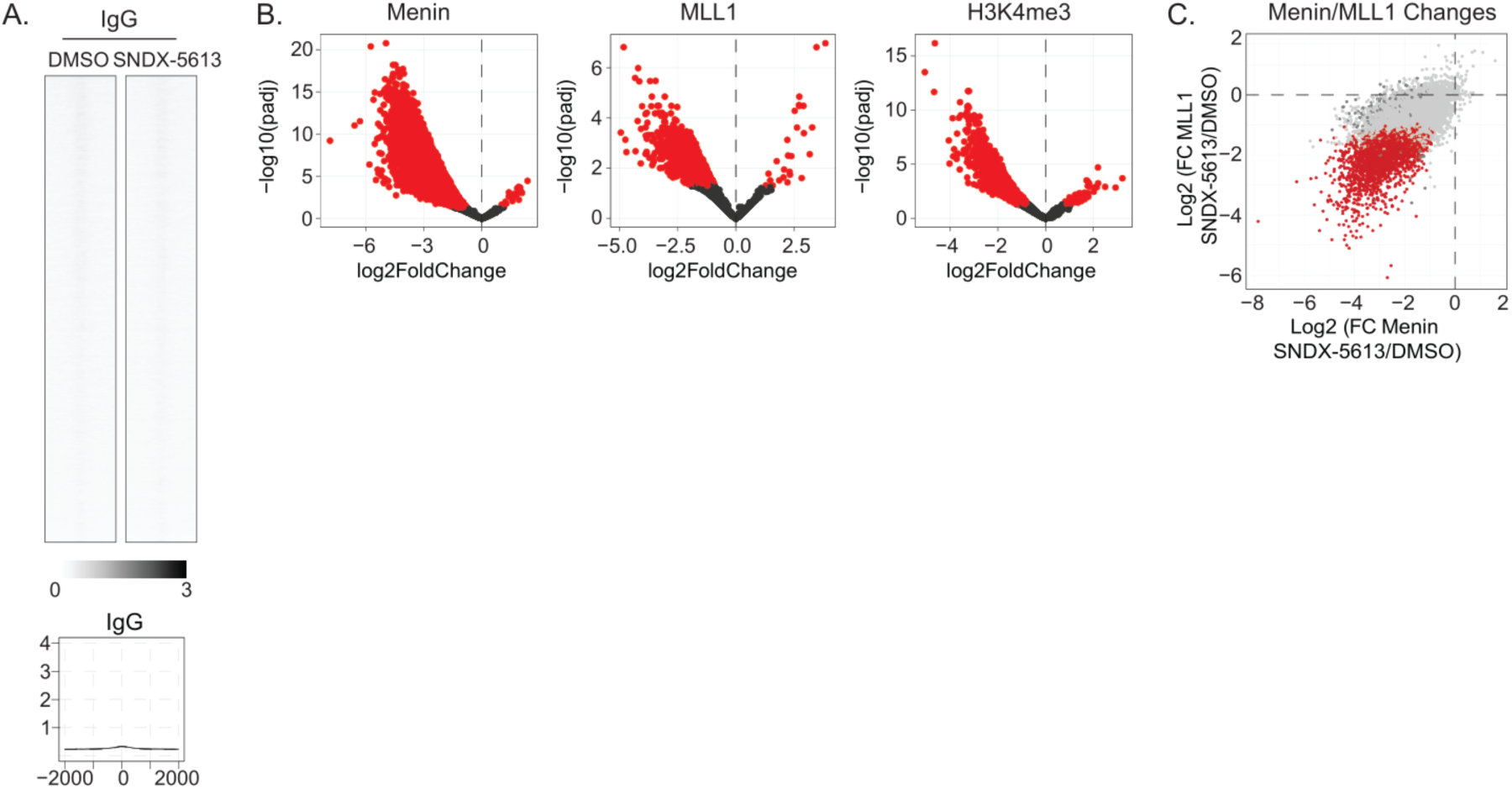
Menin inhibition leads to a global decrease in menin, MLL1, and H3K4me3 chromatin binding. (A) Heatmaps showing the correlation of promoter peaks in a ±2-kb window with occupancy of IgG across CUT&RUN-seq data from HLF cells treated with 5µM SNDX-5613 for 4 days. IgG peaks are shown for all detected menin peaks. Bottom metaplot shows the mean of overall peak signals detected at the regions comparing DMSO and SNDX-5613 treatment conditions in HLF cells. (B) Differential occupancy analysis for menin, MLL1, and H3K4me3 performed at consensus peaks for each factor. (C) Volcano plot of logarithmic differential binding of menin and MLL1 proteins in HLF cells upon SNDX-5613 treatment (5µM).

**Supplemental Figure 3.**
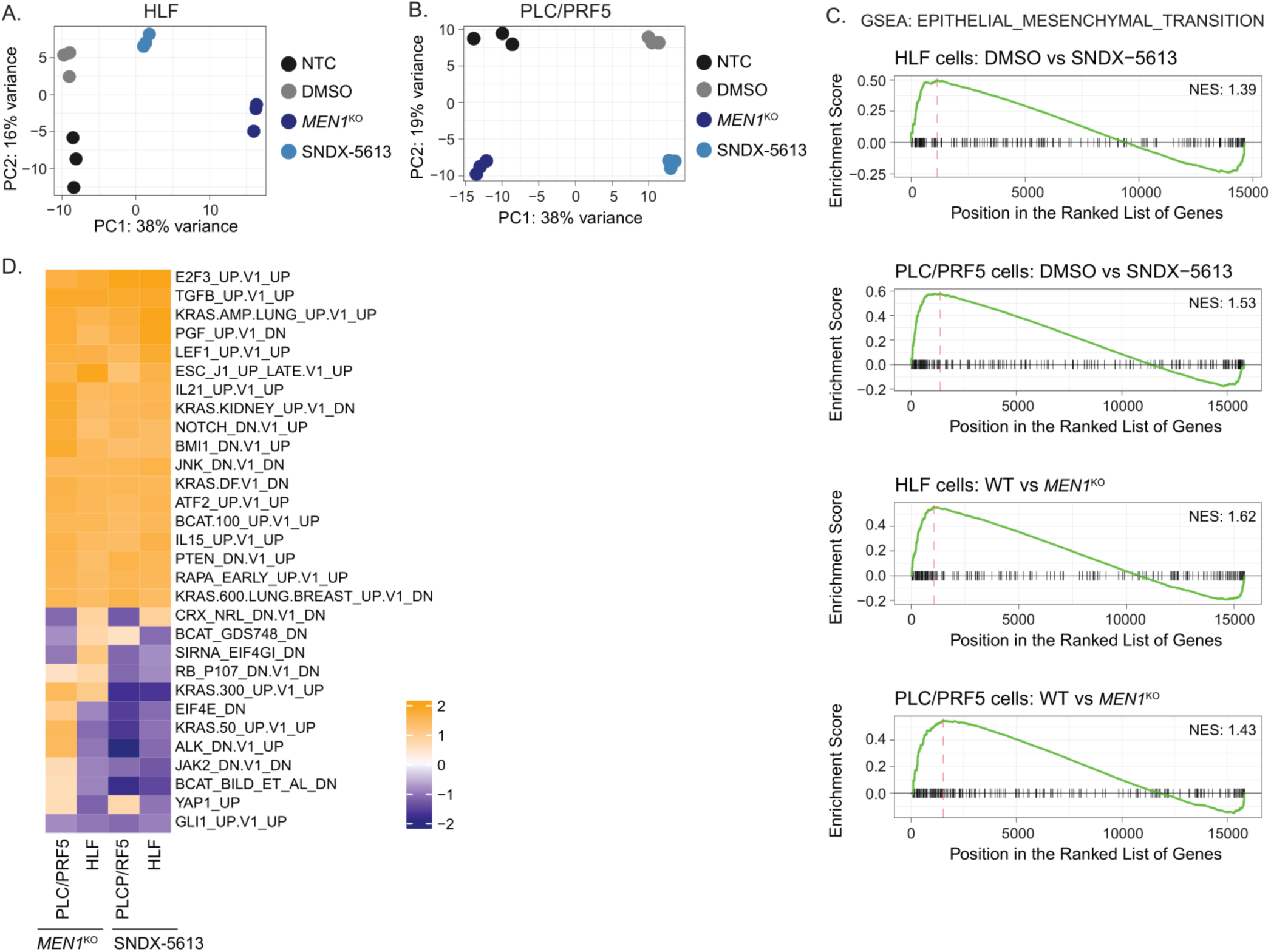
Menin inhibition alters gene expression programs. (A-B) PCA plots for replicates of HLF (A) or PLC/PRF5 (B) cells either treated with 5µM SNDX-5613 or DMSO for 4 days or carrying a *MEN1* or non-targeting gene knockout. (C) GSEA plots of gene expression changes in HLF and PLC/PRF5 cell lines both either treated with 5µM SNDX-5613 or DMSO for 4 days or carrying a *MEN1* or non-targeting gene knockout with Epithelial-Mesenchymal Transition pathway from hallmark reference gene set. (D) Comparison of GSEA ontologies with cancer gene set (C6) used as a reference affected by 5µM SNDX-5613 treatment and *MEN1* knockout in HLF and PLC/PRF5 cells.

**Supplemental Figure 4.**
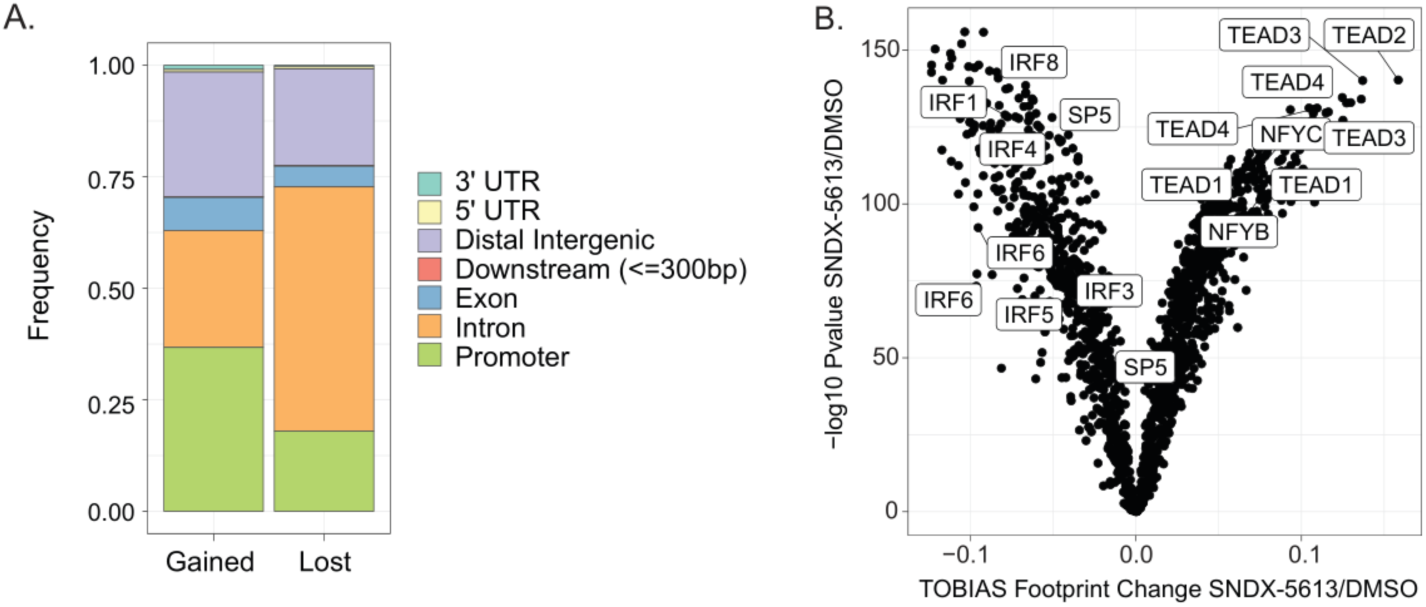
Chromatin accessibility at promoters and distal regions are disrupted by menin inhibition. (A) Genomic annotations associated with ATAC-seq peaks with increased or decreased accessibility. (B) TOBIAS analysis of motif accessibility differences in HLF cells following 5µM SNDX-5613 treatment.

**Supplemental Figure 5.**
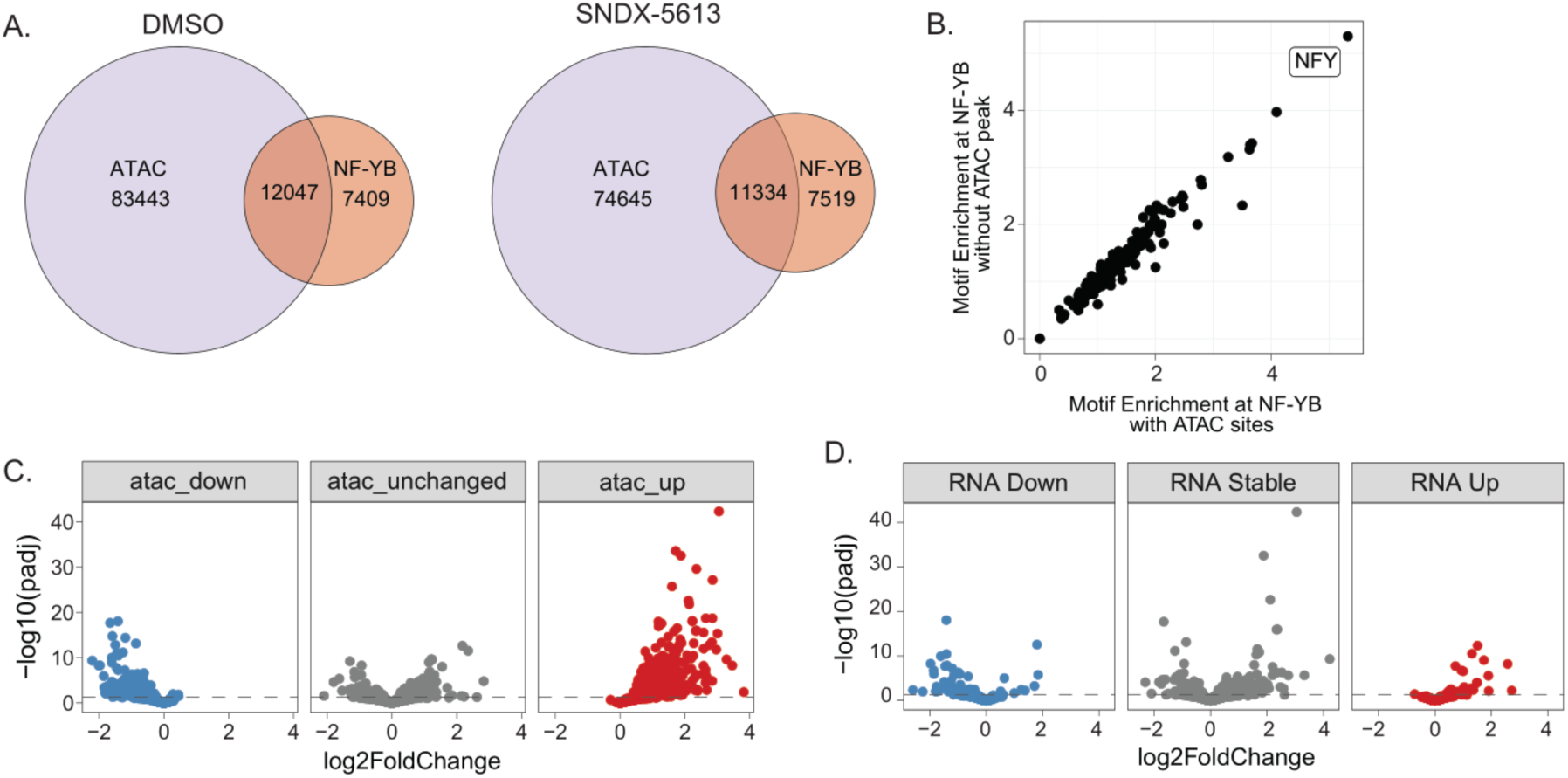
NF-YB is associated with altered chromatin accessibility at menin sites. (A). Venn diagram of overlap between NF-YB binding and ATAC-seq peaks. (B). Motif analysis using HOMER at NF-YB peaks showing the overlap between NF-YB binding peaks that either overlap (x-axis) or do not with ATAC-seq peak (y-axis). (C) Differential logarithmic occupancy of NF-YB binding at ATAC-seq peaks faceted by whether the ATAC-seq peak was upregulated, downregulated or unchanged. (D) Differential logarithmic occupancy of NF-YB binding at sites associated with expression of the nearest gene to the peak.

